# Functional diversity of Vγ9Vδ2 T cells overcomes glioblastoma state plasticity and antigen heterogeneity

**DOI:** 10.64898/2026.05.19.726243

**Authors:** P. Paris, M. Devinat, J. Ceroni, E. Guiet, J. Ollier, M. Laurent--Blond, P. Thomas, M. Gaudin, N. Joalland, A. Perrin, C.P. Delannoy, A. Rousseau, P. Menei, E. Scotet, F. Paris, C. Niaudet, S. Fougeray, S. Groux-Degroote, C. Gratas, A. Clavreul, B. Clemenceau, C. Pecqueur

## Abstract

Glioblastoma (GBM) is characterized by a high degree of cellular plasticity and intra-tumoral heterogeneity, which frequently leads to the failure of standard therapies, including immunotherapies. While chimeric antigen receptor (CAR) T cells offer a potent means of MHC-independent tumor recognition, their efficacy is hampered by the coexistence of distinct molecular states, gathered as proneural (PN) and mesenchymal (MES) phenotypes. Here, we demonstrate that gangliosides GD2 and O-acetylated GD2 (OAcGD2) are preferentially expressed by PN cells whereas MES cells display reduced expression due to upregulated ganglioside catabolism. Conversely, MES cells are known to exhibit high expression of stress-induced ligands recognized by Vγ9Vδ2 T cells. We show that while engineering Vδ2T cells with GD2- or OAcGD2-specific CAR enables the elimination of PN cells, it also facilitates the immune escape of MES cells in heterogeneous 3D-tumoroid models. Mechanistically, we reveal a hierarchy of receptor engagement, where CAR signaling predominates leading to the structural and functional exclusion of endogenous TCR from the immunological synapse. To address this receptor competition, we propose a strategy that leverages the functional effector diversity by combining untransduced and CAR-engineered Vδ2T cells. This dual approach provides a dynamic safety net by ensuring the simultaneous elimination of PN and MES cells and preventing the selective outgrowth of resistant cells. Our findings establish a conceptual framework for designing off-the-shelf immunotherapies tailored to the metabolic and phenotypic plasticity of resistant solid tumors.

## Introduction

Glioblastoma (GBM) remains one of the most lethal human cancers, with a median overall survival barely exceeding 15 months for over two decades ^1–3^. This clinical stagnation persists despite aggressive multimodal therapy, including surgery, radiation, and temozolomide, referred to as the Stupp protocol. This lack of improvement stands in stark contrast to the substantial advances in our molecular understanding of the disease, which have revealed profound intra-tumoral heterogeneity. Single-cell transcriptomic analyses have identified diverse molecular cell states that coexist within a single tumor, including oligodendrocyte progenitor-like (OPC), neural progenitor-like (NPC), and astrocyte-like (AC) states, collectively referred to as proneural (PN) states, as well as a mesenchymal-like (MES) state frequently associated with therapy resistance and metabolic plasticity ^4–7^. These states are spatially organized, with PN cells enriched in perivascular and infiltrative regions, whereas MES cells often predominate in hypoxic regions and display greater resistance to therapy ^8–13^. Moreover, standard therapies can induce a phenotypic shift toward the MES state, rendering current therapeutic approaches ineffective against this evolving cellular plasticity ^14–16^.

Immunotherapy has emerged as a promising alternative; Yet GBM poses formidable challenges to immune-based strategies, including a restrictive blood–brain barrier, limited lymphocyte trafficking, and a profound immunosuppressive tumor microenvironment ^17^, Chimeric antigen receptor (CAR) T cell trials targeting IL13Rα2, EGFRvIII, HER2, or the disialoganglioside GD2, have demonstrated manageable safety profiles and occasional transient responses ^18–23^. However, durable efficacy remains elusive, primarily due to pronounced inter- and intra-tumoral antigen heterogeneity and subsequent antigen loss under selective pressure. In particular, how engineered effectors navigate the simultaneous targeting of multiple plastic tumor states, and whether CAR-mediated signals interfere with endogenous receptor functionality, remain poorly understood.

To address this paradigm, we propose a strategy that combines CAR engineering with the versatile properties of alternative immune effectors to target the full spectrum of GBM states, rather than single antigens. We focused on human Vγ9Vδ2 T (Vδ2T) cells, as we previously demonstrated that these T cells display spontaneous cytotoxicity against MES GBM cells ^24^. This recognition is mediated by the sensing of phosphoantigen accumulation and stress-induced ligands, such as MICA/B, through the activating receptor NKG2D ^24–26^. By engineering Vδ2T cells with CAR targeting PN-enriched antigens, we aimed to create a dual recognition effector capable of covering the entire GBM spectrum. We selected GD2 and its O-acetylated derivative (OAcGD2) as CAR targets, given their high expression in tumors of neuroectodermic origin ^27^. GD2-CAR therapies have shown therapeutic potential in preclinical models ^28–31^ and promising activity in clinical trials for neuroblastoma and medulloblastoma ^32–36^. However, their use is limited by concerns regarding on-target off-tumor neurotoxicity due to GD2 expression in peripheral nerves ^32^. OAcGD2 targeting potentially offers an improved safety profile, given its absence on peripheral nerves ^27^.

In this study, we demonstrate that ganglioside expression is intrinsically linked to the metabolic state of tumor cells. We show that while CAR engineering provides potent activity against PN cells, a hierarchy of receptor engagement exists where CAR signaling outcompete the endogenous TCR, leading to the immune escape of MES cells in heterogeneous 3D contexts. Mechanistically, this is driven by the structural exclusion of TCR from CAR-driven immunological synapses. Finally, we provide a dual-recognition strategy by demonstrating that preserving functional diversity through a combination of untransduced and CAR-engineered Vδ2T cells ensures the comprehensive elimination of all tumor cells, offering a robust framework for overcoming intra-tumoral heterogeneity and state plasticity in solid tumors.

## Results

### GD2 and OAcGD2 expression correlates with GBM cell states

To assess the clinical relevance of GD2 and OAcGD2 therapeutic targets, we analyzed their expression in a cohort of 16 patients with primary and recurrent IDH-wild-type GBM from the French GBM Biobank ^37^. All patients underwent resection at recurrence following the standard Stupp protocol (Table 1). GD2 and OAcGD2 expression varied widely across tumors, ranging from approximately 10 to 90% of positive cells (Fig. 1A-C), highlighting the significant inter-patient heterogeneity. In most cases, these expression levels were maintained at recurrence. Spatially, both gangliosides exhibited restricted staining patterns, consistent with intra-tumoral heterogeneity (Fig. 1A). Although GD2 expression was consistently higher than that of OAcGD2, the two markers frequently colocalized within the same tumor region (Fig. 1A, 1D). In parallel, we evaluated the expression of antigens recognized by Vδ2 T cells, including butyrophilin 2A (BTN2A) and stress-induced ligands (ULBPs, MICs) (Supplementary Fig. 1). ULBP3 expression was significantly inversely correlated with GD2 levels (Fig. 1E). Conversely, the stress-induced ligands ULBP2/5/6 and MICA/B showed a positive correlation with each other. Altogether, these results reveal a distinct spatial organization in tumors, with regions enriched with GD2-positive cells displaying low expression of Vδ2T cell ligands, while areas with reduced GD2 expression exhibit elevated levels of these ligands.

**Fig. 1.**
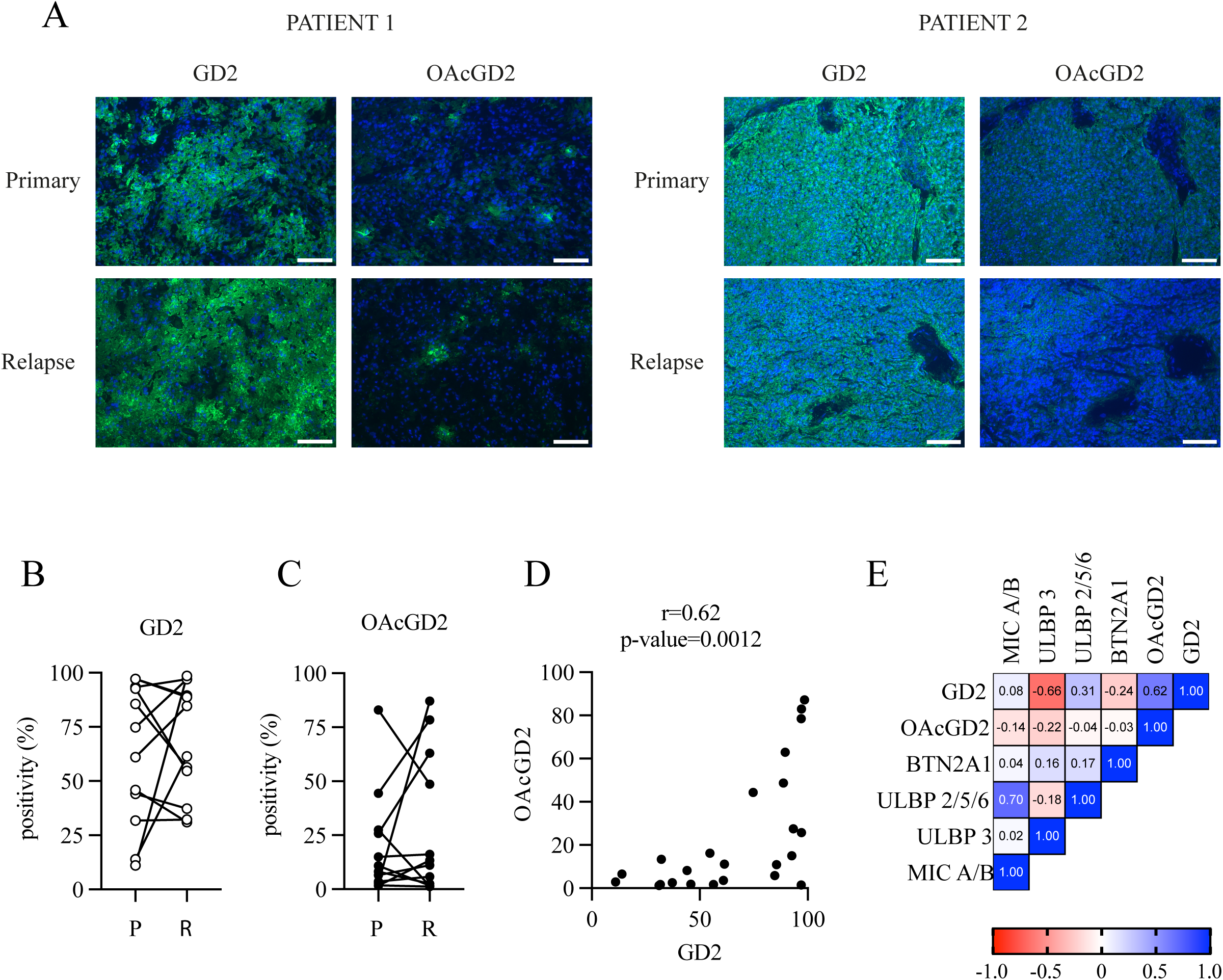
GD2 and OAcGD2 expression in GBM. A. representative images of immunofluorescence staining of GD2 and OAcGD2 on GBM samples. GD2 and OAcGD2 staining in green, DAPI in blue. The scale bar corresponds to 100 µm. B-C. Semi-quantitative analysis of GD2 and OAcGD2 in primary (P) and relapse (R) samples performed with Fiji software. D. Correlation analysis between GD2 and OAcGD2 levels. E. Correlation analysis of ganglioside expression, Vδ2 TCR ligands and NKG2D ligands across distinct tumor regions.

To further characterize this variability, we analyzed 10 patient-derived primary GBM cultures. Consistent with our tissue observations, GD2 and OAcGD2 expression was heterogeneous, positively correlated, and skewed toward higher GD2 levels (Fig. 2A-B). We then classified these cultures according to the transcriptional states defined by Neftel et al. to determine whether ganglioside expression was state-dependent ^5^. Eight cultures (GBM#9-11, 15-19) were enriched for NPC-like, OPC-like, and AC-like signatures and classified as PN, while six cultures (GBM#1-6) displayed a MES profile (Fig. 2C and Supplementary Fig. 2A). Most PN cultures exhibited higher GD2 and OAcGD2 expression than MES cells (Fig. 2D).

**Fig. 2.**
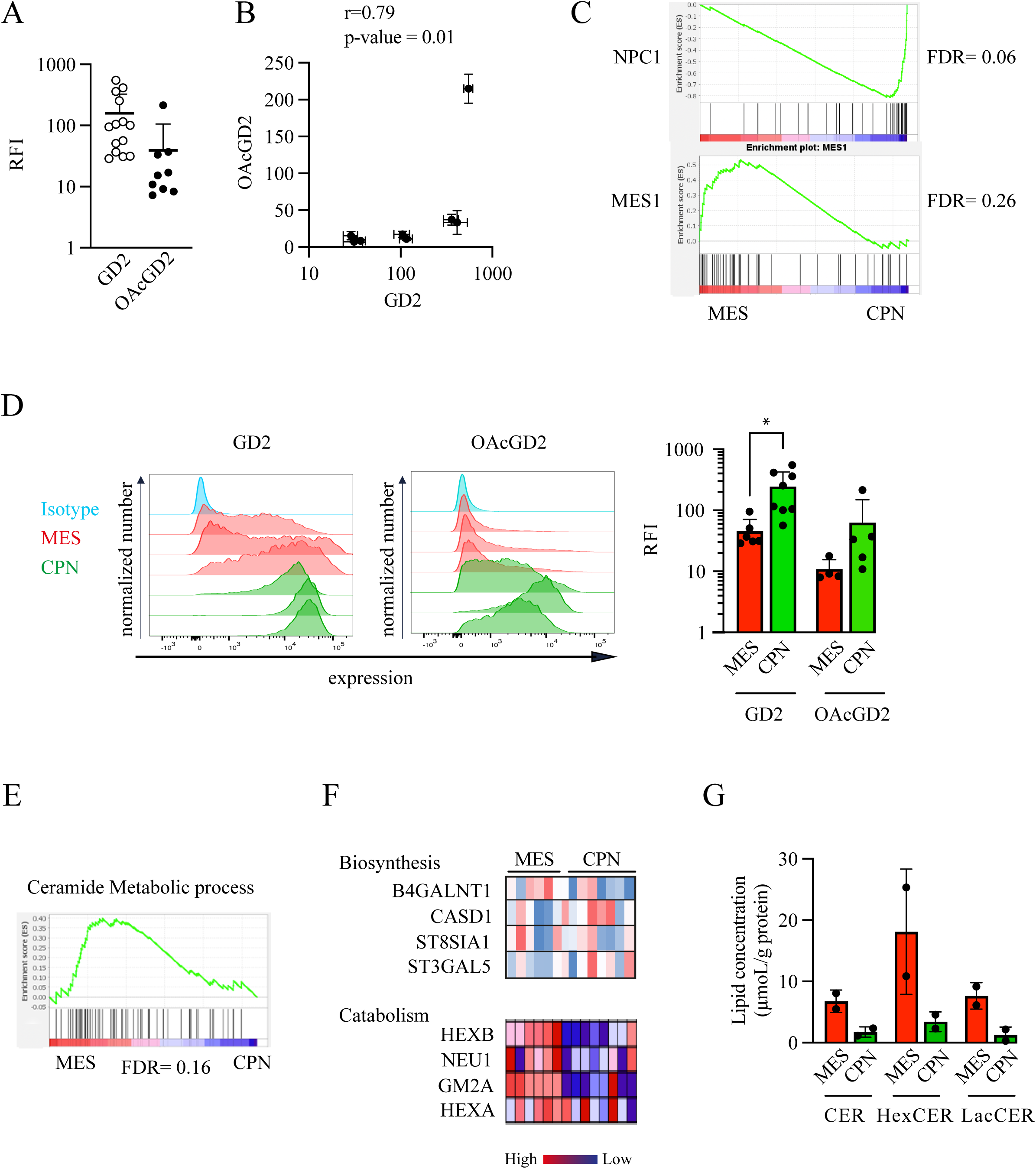
Ganglioside metabolism and expression is linked to GBM states. A-B. GD2 (A) and OAcGD2 (B) expression in 10 patient-derived primary cultures. C. Gene set enrichment analysis (GSEA) of GBM cultures into proneural (NPC1) and mesenchymal (MES1) based on metamodules defined by Neftel *et al*. D. GD2 and OAcGD2 expression based on tumor cell state. *Left panel*: representative histogram of GD2 and OAcGD2 expression in individual primary cultures; *Right panel*: Relative fluorescent intensity depending on molecular states. Each point represents an individual primary culture. E. GSEA of ceramide metabolic process in PN and MES cells. F. Differential expression of biosynthetic (B4GALNT1, CASD1, ST8SIA1, ST3GAL5) and catabolic (HEXB, NEU1, GM2A, HEXA) enzymes between cell states. G. Ceramide quantification by mass spectrometry in MES and PN cells. G. Lipidomic profiling by mass spectrometry of ceramide species in PN and MES cells.

To explore the molecular basis of this state-dependent expression, we performed gene set enrichment analyses (GSEA) focusing on sphingolipid metabolism. Surprisingly, MES cells, and not PN cells, were enriched in signatures associated with ceramide metabolism, glycosphingolipid biosynthesis, and ganglioside metabolism (Fig. 2E, Supplementary Fig. 2B-C). However, the expression of key biosynthetic enzymes, such as B4GALNT1, ST8SIA1, ST3GAL5, and CASD1, remained largely unchanged across cell states (Fig. 2F). In contrast, genes involved in ganglioside degradation, such as *HEXB*, *GM2A*, and *NEU1*, were significantly upregulated in MES cells, but weakly expressed in PN cells. Supporting these transcriptomic findings, mass spectrometry revealed higher levels of ceramide, hexosylceramide, and lactosylceramide in MES cells, along with a more complex and diverse ganglioside profile, compared to the simpler profile observed in PN cells (Fig. 2G and Supplementary Fig. 2D). In contrast, PN GBM15 cells exhibited a simpler ganglioside profile with fewer species detected.

Altogether, these results demonstrate that while both GD2 and OAcGD2 are expressed by GBM cells, their expression is modulated by tumor metabolic and molecular states. The enrichment of gangliosides in PN cells provides a compelling biological rationale for combining CAR-mediated targeting with the innate cytotoxicity of Vδ2T cells.

### CAR-engineering confers potent and antigen-specific cytotoxicity against PN cells

To selectively target PN cells, we engineered human Vδ2T cells with second-generation CAR directed against either GD2 or OAcGD2. Truncated variants lacking the CD3ζ signaling domain (ΔCAR) were used as negative controls (Fig. 3A). Following transduction and selection, we first evaluated CAR-dependent activation against PN cells, which express high surface levels of both gangliosides. As expected, untransduced (UT) and ΔCAR Vδ2T cells showed minimal degranulation in the presence of PN cells (Fig. 3B). In contrast, GD2-ζCAR Vδ2T cells exhibited robust and specific CD107a surface expression across all tested PN cultures. OAcGD2-ζCAR Vδ2T cells were selectively activated by PN cells with high OAcGD2 expression (GBM#10 and GBM#15), suggesting that OAcGD2-ζCAR activation is sensitive to the antigen threshold. Consistent with these activation patterns, both GD2-ζCAR and OAcGD2-ζCAR Vδ2T cells secreted high levels of IFN-γ and TNF-α upon co-culture with GBM#15 cells (Fig. 3C). We noted that OAcGD2-ζCAR Vδ2T cells produced detectable IFN-γ in the absence of targets, suggesting a baseline level of tonic signaling (Supplementary Fig. 3A-C). To formally confirm the antigen specificity of ζCAR constructs, we genetically silenced B4GALNT1 and CASD1, key enzymes involved in GD2 and OAcGD2 biosynthesis, respectively. Silencing B4GALNT1 in PN cells significantly reduced GD2 expression and subsequent GD2-ζCAR Vδ2T degranulation, whereas UT and ΔCAR T cells were unaffected (Fig 3D-E, Supplementary Fig. 3D). The residual degranulation observed in GD2-ζCAR Vδ2T cells likely reflects either residual GD2 expression or potential cooperativity between ζCAR and Vδ2 T cell recognition. Silencing CASD1 resulted in a complete loss of OAcGD2 expression (Fig. 3F), and a concomitant reduction in GD2 levels (Supplementary Fig. 3E). CASD1 silencing abolished OAcGD2-ζCAR Vδ2T degranulation (Fig. 3G), confirming the dependency on OAcGD2.

**Fig. 3.**
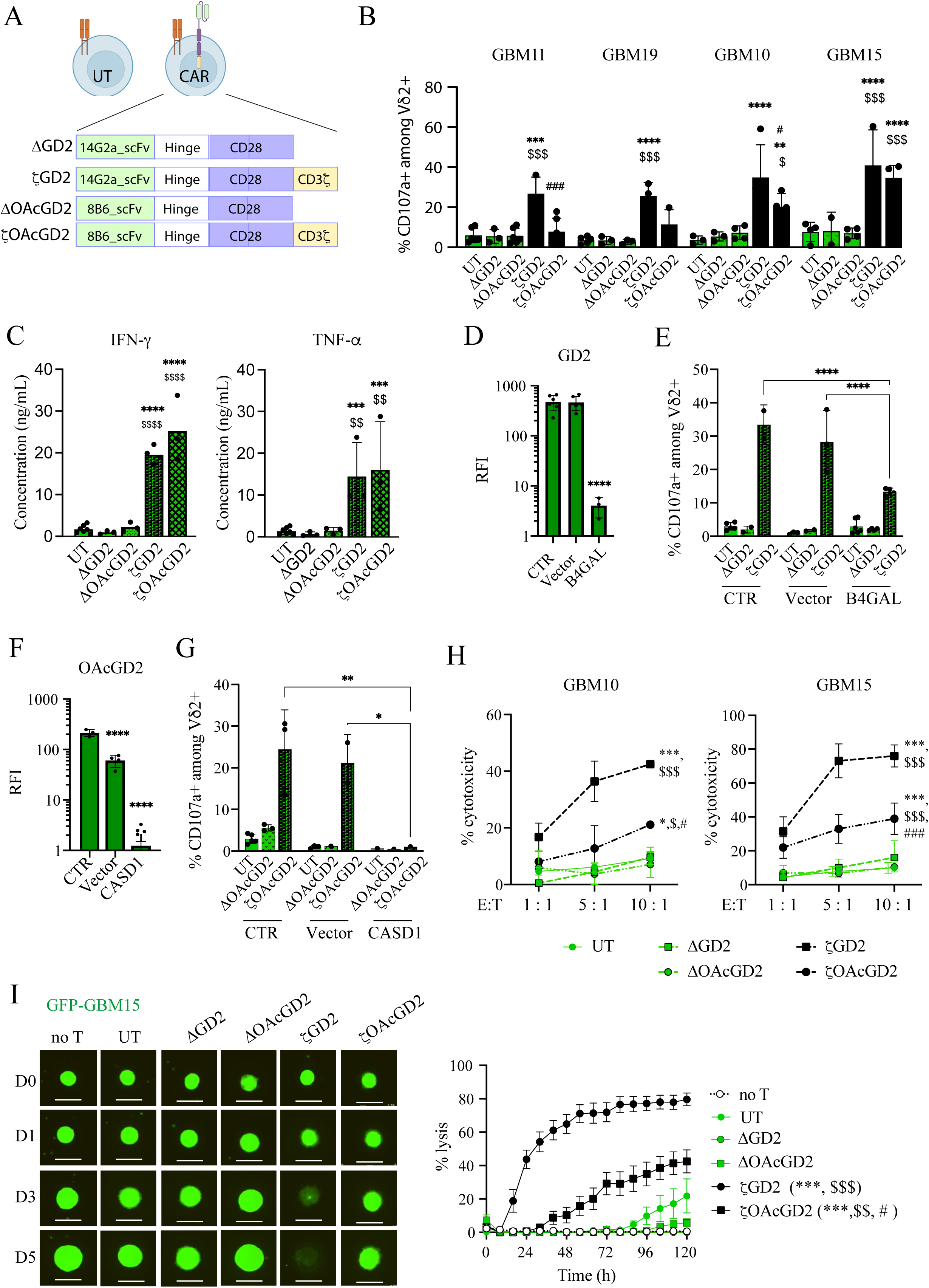
CAR transduction allows PN killing by Vδ2T cells. A. Schematic representation of second-generation ζCAR constructs and truncated ΔCAR controls. B. Degranulation of Vδ2T cell subsets following coculture with PN cells (E:T ratio of 1:1). C. Secretion of IFNγ (left) and TNFα (right) by Vδ2T cell subsets following coculture with PN cells. D. GD2 expression following B4GALNT1 silencing in PN cells. E. Degranulation of Vδ2T cell subsets following coculture with PN cells after B4GALNT1 silencing. F. OAcGD2 expression following CASD1 silencing in PN cells. G. Degranulation of Vδ2T cell subsets following coculture with PN cells after CASD1 silencing. H. Short-term cytotoxicity (24 hours) of Vδ2T cell subsets following coculture with PN cells (*left panel* GBM15 cells; *right panel* GBM10 cells). I. Long-term cytotoxicity (5 days) of Vδ2T cells following coculture with PN cells (E:T ratio of 10:1). *Left panel*: representative images of 3D-tumoroid (scale bar=0.8mm); *right panel*: Lysis quantification over time (%). Statistical significance was determined using two-way ANOVA with Tukey’s post-hoc test. Asterisks (*) denote significant differences compared to the UT group, dollar signs ($) indicate significant differences between ζCAR and corresponding ΔCAR, hash sign (#) indicate significant differences between GD2-ζCAR and OAcGD2-ζCAR.

Next, we assessed the cytotoxic potential of ζCAR Vδ2T in 24-hour co-culture assays at varying effector-to-target (E:T) ratios (Fig. 3H). Both GD2-ζCAR and OAcGD2-ζCAR Vδ2T cells significantly killed PN tumor cells in an E:T-dependent manner. Overall, GD2-ζCAR Vδ2T exhibited the highest cytotoxic potential. UT and ΔCAR Vδ2T cells displayed no significant cytotoxicity against PN cells. To evaluate these effectors in a more complex context, we performed longitudinal live cell imaging of GFP-expressing PN-derived 3D-tumoroids over five days. Both ζCAR-engineered Vδ2T cells induced a progressive elimination in a CAR- and E:T-dependent manner (Fig. 3I and Supplementary Fig. 3F-G).

Collectively, these results establish that CAR engineering successfully redirects Vδ2T cells to eliminate PN cells through potent antigen-specific cytotoxicity.

### CAR engineering preserves Vδ2T cell cytotoxicity against MES cells

We explore whether CAR expression alters Vδ2 T cell ability to recognize and eliminate MES cells. As expected, UT Vδ2T cells exhibited significant degranulation in the presence of MES cells, but remained unresponsive to PN cells (Fig. 4A). Importantly, CAR transduction did not impair Vδ2T cell degranulation activity across MES models (Fig. 4B). In some instances, such as GBM#1 and GBM#4, CAR-engineered cells showed enhanced degranulation compared to UT cells. Regarding cytokine production, both GD2-ζCAR and OAcGD2-ζCAR Vδ2T cells secreted higher absolute levels of IFN-γ and TNF-α upon co-culture with MES cells than their UT and ΔCAR counterparts (Fig. 4C). However, when accounting for the higher baseline cytokine production observed in CAR-engineered cells, the relative fold induction following tumor cell exposure remained comparable across all Vδ2T cell subsets (Supplementary Fig. 4A). We then evaluated the cytotoxic function in both short-term (24 h) and long-term (5 d) assays. In 24-hour co-cultures, all Vδ2T cell subsets, regardless of CAR expression, effectively killed MES tumor cells in an E:T-dependent manner (Fig. 4D). Interestingly, CAR engineering did not further enhance the cytotoxicity against MES cells, as compared to UT cells. These findings were confirmed using RFP-expressing MES-derived 3D-tumoroids. Longitudinal live-cell imaging demonstrated that all Vδ2T cell subsets displayed similar, robust killing over five days (Fig. 4E, Supplementary Fig. 4B-C).

**Fig. 4.**
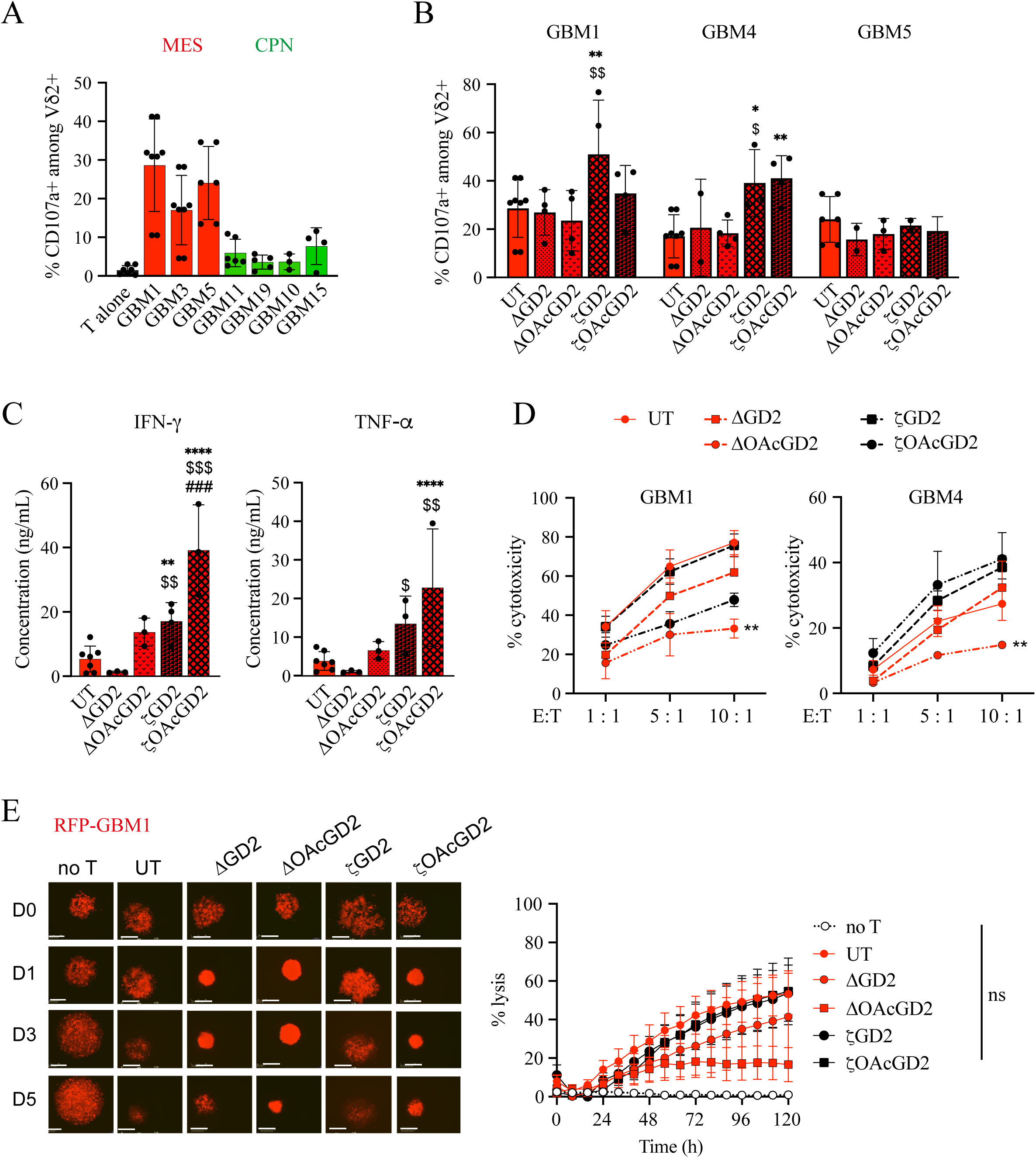
CAR transduction does not alter Vδ2T cytotoxicity against MES cells. A. Degranulation of UT Vδ2T cells in presence of MES and PN cells. B. Degranulation of Vδ2T cell subsets following coculture with MES cells (E:T ratio of 1:1). C. Secretion of IFNγ (left) and TNFα (right) by Vδ2T cell subsets following coculture with MES cells. D. Short-term cytotoxicity (24 hours) of Vδ2T cell subsets following coculture with MES cells. E. Long-term cytotoxicity (5 days) of Vδ2T cells following coculture with MES cells (E:T ratio of 10:1). *Left panel*: representative images of 3D-tumoroid (scale bar=0.8mm); *right panel*: Lysis quantification over time (%). Statistical significance was determined using two-way ANOVA with Tukey’s post-hoc test. Asterisks (*) denote significant differences compared to the UT group, dollar signs ($) indicate significant differences between ζCAR and corresponding ΔCAR, hash sign (#) indicate significant differences between GD2-ζCAR and OAcGD2-ζCAR.

Collectively, these results demonstrate that CAR engineering preserves Vδ2T cell cytotoxicity, ensuring that they remain effective against MES cells while gaining new specificity for PN cells.

### Intra-tumoral heterogeneity restricts the dual-targeting of ζCAR Vδ2T cells in 3D-tumoroids

Given that MES and PN cells coexist within individual tumors, we investigated the efficacy of GD2-ζCAR Vδ2T cells in a mixed-state context. We first assessed short-term cytotoxicity in 2D co-cultures composed of an equal ratio of RFP-labeled MES cells and GFP-labeled PN cells (Fig. 5A). Across all conditions, tumor cell killing increased proportionally with E:T ratio across all conditions (Fig. 5B). GD2-ζCAR Vδ2T cells exhibited superior overall cytotoxicity at all ratios. Notably, at the lowest E:T ratio (1:1), UT Vδ2T cells mediated only 15% tumor cell killing, whereas GD2-ζCAR Vδ2T cells induced over 40% tumor cell killing. GD2-ΔCAR T cells displayed intermediate cytotoxicity (29%), suggesting that ΔCAR may facilitate TCR-mediated cytotoxicity, perhaps by favoring effector-target contact. To decipher the state-specific effects, we independently analyzed MES and PN cell killing. Under these 2D conditions, MES cell killing remained similar across all T cell subsets (Fig 5C). In contrast, GD2-ζCAR T cells were highly effective in killing PN cells, while UT Vδ2T cells had minimal impact (Fig 5D). Again, GD2-ΔCAR T cells exhibited intermediate levels of PN killing, further suggesting that the ΔCAR-antigen interaction might indirectly potentiate TCR-mediated recognition by favoring cell-cell contact.

**Fig. 5.**
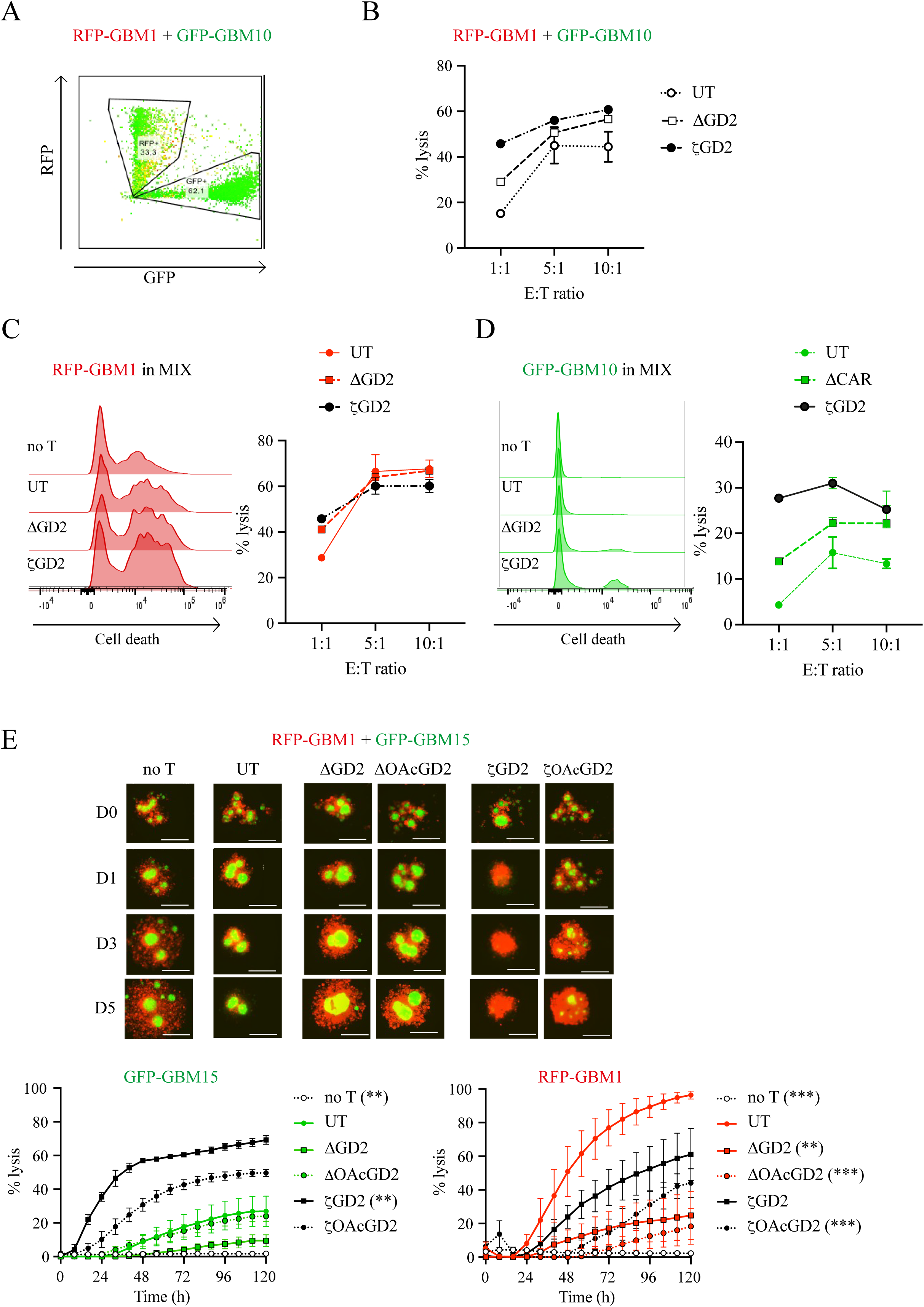
MES cells escape CAR Vδ2T cell killing in mixed tumoroids. A. Representative FACS analysis of 2D-mixed cultures with equal ratios of RFP-MES and GFP-PN cells (1:1). B-D. Short-term cytotoxicity (24 hours) of Vδ2T cell subsets against 2D-mixed cultures (B), RFP-MES cells (C) and GFP-PN cells (D) within mixed cultures. E. Long-term cytotoxicity (5 days) of Vδ2T cell subsets against mixed 3D-tumoroids (E:T ratio of 10:1). *Top panel*: representative tumoroid images over time (scale bar=0.8mm); *bottom panel*: % lysis of GFP- and RFP-expressing tumor cells over time. Statistical significance was determined using two-way ANOVA with Tukey’s post-hoc test. Asterisks (*) denote significant differences compared to the UT group.

We expanded our experiments to include mixed 3D-tumoroid models composed of equal proportions of MES (RFP-GBM1) and PN cells (GFP-GBM15). Strikingly, in this heterogeneous 3D context, distinct dynamics emerged, leading to MES cell immune escape (Fig. 5E). As expected, UT Vδ2T cells preferentially eliminated MES cells while sparing PN cells. Conversely, GD2-ζCAR and OAcGD2-ζCAR Vδ2T cells efficiently targeted and killed the PN cells. However, neither GD2-ζCAR nor OAcGD2-ζCAR Vδ2T cells achieved efficient elimination of MES cell in this 3D setting, contrasting with our 2D findings. Similar results were observed in a second mixed 3D-model (Supplementary Fig. 5).

These findings indicate that within the complex 3D GBM context, CAR-engineering may prioritize the elimination of high-antigen PN cells at the expense of MES cell killing, ultimately failing to provide simultaneous dual coverage.

### CAR signaling dominates the immunological synapse and outcompetes endogenous TCR recognition

To elucidate the mechanisms underlying MES cell escape in mixed tumoroids, we investigated the relative contributions of CAR and TCR signaling using genetic silencing and pharmacological inhibition. We used a BTN3 inhibitor (BTN3i) to block Vδ2 TCR-mediated recognition and B4GALNT1 silencing to abolish GD2 expression in MES cells (Fig. 6A). While B4GALNT1 silencing had no effect on UT Vδ2T cell activation, it significantly reduced GD2-ζCAR Vδ2T cell degranulation (Fig. 6B). Conversely, BTN3i completely prevented UT Vδ2T cell activation but did not affect GD2-ζCAR T cell activation. Interestingly, while neither GD2 loss nor BTN3i alone significantly affected GD2-ΔCAR Vδ2T cells, simultaneous inhibition of both pathways was required to reduce their degranulation, further supporting the hypothesis that ΔCAR facilitates TCR-dependent interaction (Fig. 6B).

**Fig. 6.**
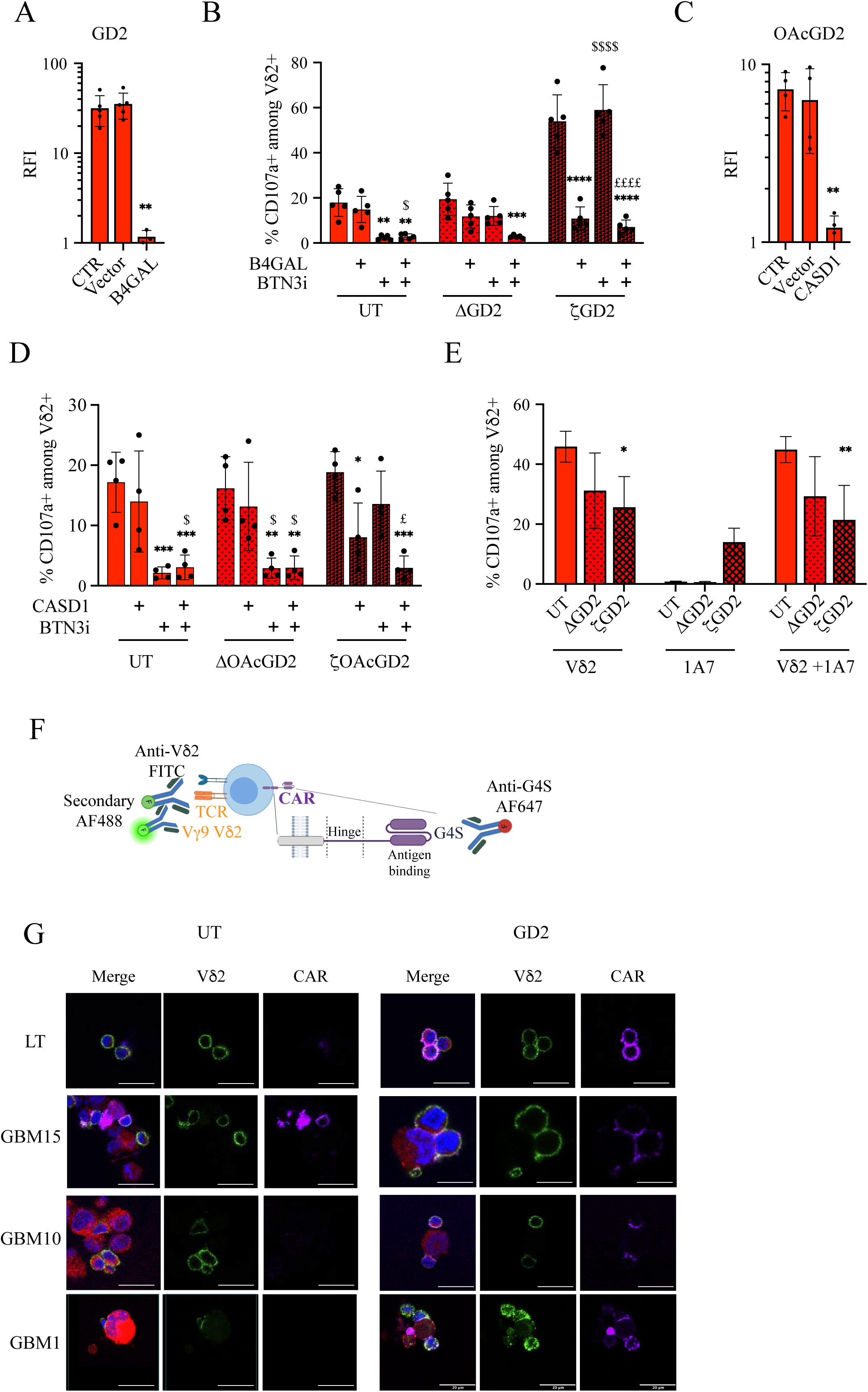
CAR signaling predominates over TCR recognition. A. GD2 expression following B4GALNT1 silencing in MES cells (GBM#1). B. Degranulation of Vδ2T cell subsets following coculture with MES cells after B4GALNT1 silencing and/or addition of BTN3 inhibitors (BTN3i). C. OAcGD2 expression following CASD1 silencing in MES cells. D. Degranulation of Vδ2T cell subsets following coculture with PN cells after CASD1 silencing. E. Degranulation assay under single or dual stimulation with either TCR agonist and 1A7 anti-idiotype CAR-mimetic. F. Schematic representation of synapse characterization. G. Representative confocal images of the immunological synapse. Statistical significance was determined using two-way ANOVA with Tukey’s post-hoc test. Asterisks (*) denote significant differences compared to the UT group, dollar signs ($) indicate significant differences between ζCAR and corresponding ΔCAR, hash sign (#) indicate significant differences between GD2-ζCAR and OAcGD2-ζCAR.

Similar results were obtained for OAcGD2-ζCAR T cells. CASD1 silencing significantly reduced OAcGD2 expression in MES cells without altering GD2 levels (Fig. 6C and Supplementary Fig. 6). While UT cell activation relied strictly on Vδ2 TCR recognition, OAcGD2-ζCAR activation was driven by OAcGD2 expression (Fig. 6D). However, unlike GD2-ζCAR T cells, full inhibition of OAcGD2-ζCAR T cells required simultaneous blockade of both TCR and CAR-mediated pathways, suggesting a degree of cooperation when the CAR antigen is less abundant. To further explore this, we stimulated Vδ2T subsets with either a Vδ2 TCR agonist, an anti-idiotype antibody (1A7) mimicking GD2 engagement, or both (Fig. 6E). Although both stimuli independently triggered strong and specific degranulation, dual stimulation did not enhance activation beyond the levels achieved by TCR stimulation alone, indicating a lack of additive signaling.

Finally, we analyzed the spatial organization of the immunological synapse (Fig. 6F). Under resting conditions, both the Vδ2 TCR and GD2-ζCAR were homogeneously distributed on the T cell surface (Fig. 6G). Upon co-culture with PN cells, UT Vδ2T cells failed to form synapses, whereas GD2-ζCAR T cells displayed robust CAR polarization at the effector-target interface. Strikingly, the endogenous TCR was excluded from CAR-driven synapses. When challenged with MES cells, UT Vδ2 T cells showed the expected TCR mobilization. However, in GD2-ζCAR T cells, the CAR was preferentially recruited at the synapse over the Vδ2 TCR.

Altogether, these results reveal that ζCAR reshapes the immunological synapse, leading to a functional predominance of CAR signaling when the antigen is present. This hierarchical organization explains why CAR engagement can outcompete and effectively blind the endogenous TCR-mediated response against MES cells in heterogeneous environments.

### Combining UT and CAR-engineered Vδ2T cells overcome tumor heterogeneity in 3D models

Given that ζCAR expression restricts the innate targeting of MES cells in a mixed environment, we evaluated whether a combinatorial approach mixing UT with ζCAR-engineered Vδ2T cells could overcome GBM state heterogeneity. We performed longitudinal killing assays in mixed 3D-tumoroids using a 1:1 combination of UT and ζCAR Vδ2T cells. Consistent with our previous observations, UT Vδ2T cells selectively killed MES cells while sparing PN cells, whereas GD2-ζCAR T cells preferentially targeted PN cells (Fig. 7A). Strikingly, the combination of UT and GD2-ζCAR T cells resulted in robust and simultaneous elimination of both tumor subpopulations, even at lower E:T ratios (Fig. 7A-C). Similar results were observed in the second mixed 3D-model (Supplementary Fig. 7A). Detailed kinetic analysis revealed a distinct and sequential pattern of tumor elimination with rapid elimination of PN cells at early time points by GD2-ζCAR T cells, followed by steady, delayed killing of MES cell by UT Vδ2T cells. Interestingly, this synergistic effect was only observable in long-term 3D assays as short-term 2D killing assays failed to show any significant benefit of combining UT and GD2-ζCAR T cells over GD2-ζCAR T cells alone (Supplementary Fig. 7).

**Fig. 7.**
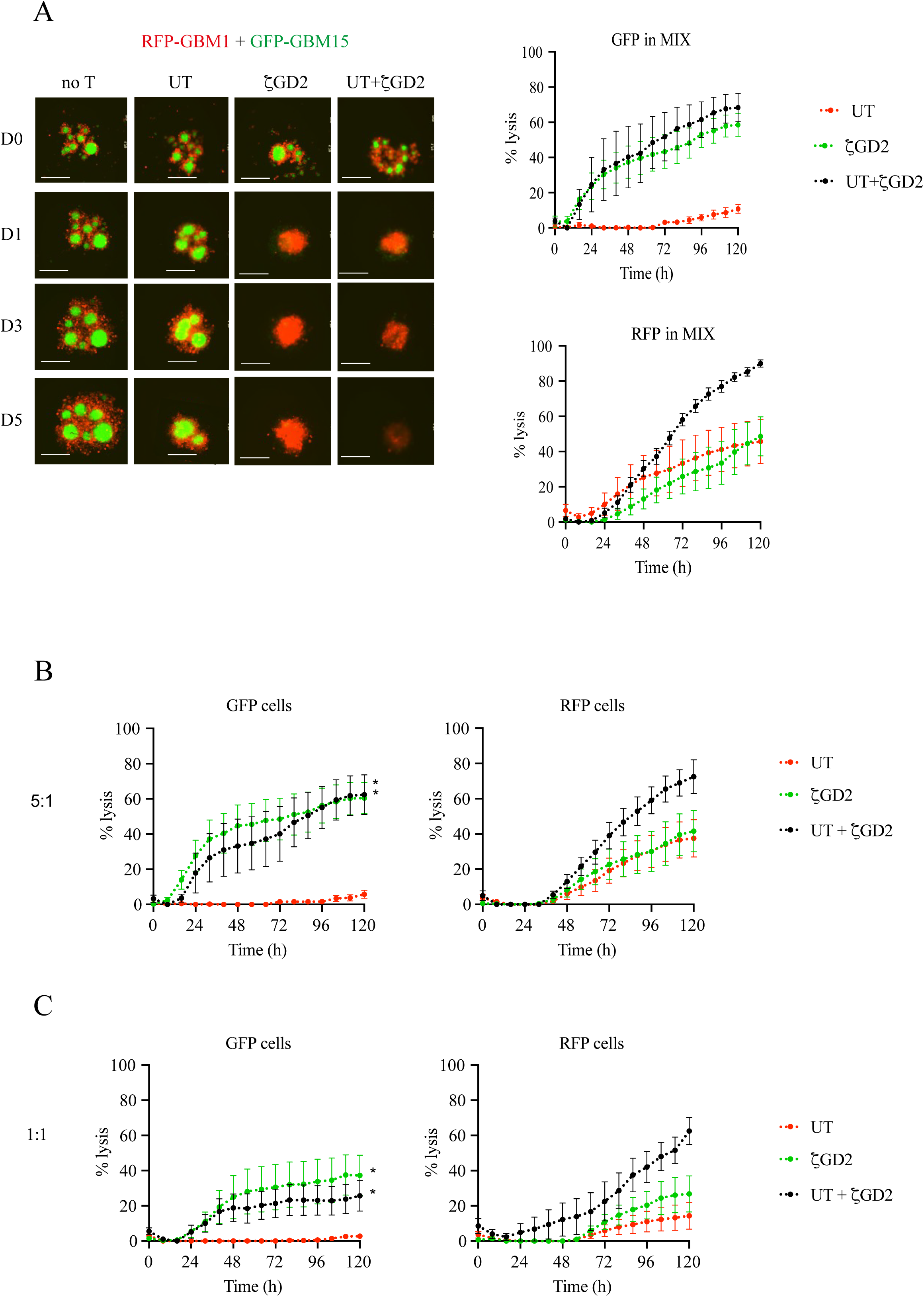
GD2-CAR Vδ2T cells require UT cell support to ensure coordinated heterogeneous tumor clearance. A-C. Long-term cytotoxicity (5 days) of Vδ2T cell subsets against mixed 3D-tumoroids at E:T ratio of 10:1 (A), 5:1 (B) and 1:1 (C).

Collectively, these findings demonstrate that a heterogeneous effector strategy, leveraging both engineered and innate recognition, can effectively mitigate intra-tumoral state plasticity, offering a promising therapeutic framework for GBM treatment.

## Discussion

Intra-tumoral heterogeneity, spanning both molecular profiles and functional states, remains the primary obstacle to efficient immunotherapy in GBM. In this study, we demonstrate that the therapeutic targets of GD2 and OAcGD2 are heterogeneously distributed across GBM cell states, with preferential enrichment in the PN state. Conversely, ligands associated with Vδ2 TCR activation are enriched in the MES state^24^. While CAR-engineering redirects Vδ2T cells to eliminate both states when evaluated independently, we uncover a critical hierarchy of receptor engagement. Indeed, ζCAR signaling predominates over endogenous TCR. In 3D models mimicking tumor heterogeneity, this dominance leads to immune escape of MES cells. Importantly, we show that preserving the functional diversity of the Vδ2T cell product by combining UT and CAR-engineered T cells restores comprehensive tumor elimination.

A key mechanistic insight from our study is the link between tumor state and ganglioside metabolism. Metabolic rewiring is a hallmark of GBM states ^7,13,38^, and our data suggest that it directly dictates the ganglioside landscape ^39,40^. While PN cells maintain high GD2 and OAcGD2 levels, MES cells exhibit a more complex ceramide and ganglioside profile, driven by the upregulation of catabolic enzymes such as HEXB and NEU1 ^41^. This state-dependent antigen distribution, coupled with the differential expression of stress-induced ligands, creates a divergent susceptibility to immune effectors. As tumor plasticity is increasingly recognized as a driver of resistance ^5,42^, single-antigen therapies may inadvertently select resistant states. Our 3D-tumoroid findings underscore that effective GBM immunotherapies must account for these dynamic shifts rather than targeting a static antigenic profile ^43–45^.

Our study revealed that ζCAR expression reshapes Vδ2T cells, both functionally and structurally. Structurally, we observed the exclusion of endogenous TCR from the immunological synapse upon CAR engagement. This physical displacement is likely compounded by a signaling trade-off within the effector cell. Indeed, the high-intensity signal delivered by the CD3ζ-based CAR may sequester critical downstream molecules, such as ZAP70 and LAT, effectively monopolizing the intracellular machinery at the expense of the TCR ^46^. This hierarchical dominance ensures the elimination of antigen-high PN cells but blinds the effector to antigen-low MES cells when both are present. Interestingly, the intermediate phenotype observed with ΔCAR constructs suggests that the physical presence of CAR may modulate the activation threshold by stabilizing effector-target contacts, even in the absence of canonical signaling.

To overcome this receptor competition, we leveraged a mixture of UT- and CAR-engineered T cells, achieving complete coverage by combining TCR-driven recognition of MES cells with CAR-mediated specificity for PN cells. This dual strategy is particularly well-suited to address the high phenotypic plasticity of GBM cells. Beyond pre-existing heterogeneity, PN cells frequently undergo mesenchymal transition under therapeutic stress ^15,47^. By deploying a heterogeneous effector population, we create a dynamic safety net since if a PN cell escapes CAR-mediated pressure by transitioning to a MES-like state or losing its antigen, it remains susceptible to the innate TCR-mediated cytotoxicity of UT cells. This approach ensures that the therapy remains as adaptive as the tumor itself, preventing the selective outgrowth of resistant cells driven by state transition. Furthermore, our strategy is positioned to synergize with the current standard of care, which is known to upregulate stress-induced ligands expression ^48–52^. By combining CAR specificity with the innate ability of Vδ2T cells to sense treatment-induced cell stress, we further strengthen the clinical potential of ζCAR Vδ2T cells in the context of early line intervention.

Beyond biological efficacy, this strategy offers significant translational and manufacturing advantages ^53,54^. Retaining UT cells within the final product bypasses the need for stringent selection, preserves the overall fitness of the effector population, and improves feasibility in autologous settings, where the expansion potential of T cells may be limited ^34,54^. Our findings challenge the current paradigm in CAR T cell manufacturing, which typically strives for the maximum purity. Instead, we propose that an optimal therapeutic product may be defined by a ratio of engineered and native effectors, maximizing multi-antigenic coverage without losing the versatile, innate sensing abilities of Vδ2T cells ^23,55–59^.

In conclusion, by integrating tumor state biology, ganglioside metabolism, and the hierarchy of receptor engagement, our study establishes a conceptual framework for designing immunotherapies tailored to GBM plasticity. This model of functional diversity provides a roadmap to overcome the antigen dilemma and immune escape, with potential applications in other resistant solid tumors.

## Acknowledgments

We thank OGD2 Pharma and ImCheck Therapeutics for providing OGD2 and 103.2 BTN3 antibodies, respectively. We are grateful to J. Amiaud and C. Charrier for immunohistochemistry support, and to Odile Blanchet and Morgane Dhondt for assistance with biobanking and sample logistics at the Angers Biological Resource Center (CRB_BB-0033-00038). We acknowledge the contribution of the MicroPicell imaging core facility, the UTE animal facility, and the Cytocell cytometry platform. We are thankful to the Labex IGO program for support. We also thank the coordinators of the French Glioblastoma Biobank (FGB), Prof. Philippe Menei (CHU Angers), Prof. Dominique Figarella-Branger (APHM Marseille), and Dr. Luc Bauchet (CHU Montpellier), as well as the clinical and technical personnel involved. This work was supported by the Institut National du Cancer (INCa), Ligue Nationale Contre le Cancer (LNCC), and Fondation ARC through the *Programme d’Actions Intégrées de Recherche (PAIR) – Gliome* initiative.

## Supplementary Figures

**Supplementary Fig. 1 related to Fig. 1.** Representative images of immunohistochemistry staining of ligands recognized by Vδ2T cells on patient samples. The scale bar corresponds to 100 µm.

**Supplementary Fig. 2 related to Fig. 2.** A-C. Gene Set Enrichment Analysis (GSEA) of primary cultures based on NPC2, AC, MES2 and OPC defined by Neftel (A), Glycosphingolipid biosynthesis (B) and Ganglioside metabolism (C). D. Mass spectrometry analysis of ganglioside profile in MES and PN cells.

**Supplementary Fig. 3 related to Fig. 3.** A-C. Vδ2T cell subset activation in the absence of tumor cells through degranulation assay (A), and secretion of IFNγ (B) and TNFα (C). D. Mass spectrometry analysis of ganglioside profile in MES and PN cells after B4GALNT1 silencing. E. GD2 expression following CASD1 silencing in PN cells. F. Long-term cytotoxicity (5 days) of Vδ2T cell subsets against 3D-tumoroids of GBM10 cells (PN) (E:T ratio of 10:1). G. Long-term cytotoxicity (5 days) of GD2-ζCAR Vδ2T against 3D-tumoroids of GBM15 cells (PN) at various E:T ratios. Statistical significance was determined using two-way ANOVA with Tukey’s post-hoc test. Asterisks (*) denote significant differences compared to the UT group, dollar signs ($) indicate significant differences between ζCAR and corresponding ΔCAR, hash sign (#) indicate significant differences between GD2-ζCAR and OAcGD2-ζCAR.

**Supplementary Fig. 4 related to Fig. 4.** A. Cytokine secretion in presence tumor cells over in absence of tumor cells. B-C. Long-term cytotoxicity (5 days) of UT (B) and GD2-ζCAR (C) Vδ2T cells against 3D-tumoroids of GBM1 cells (MES) at various E:T ratios.

Supplementary Fig. 5 related to Fig. 5. Long-term cytotoxicity (5 days) of UT, GD2-ζCAR against mixed tumoroids composed of RFP-GBM4:GFP-GBM10 (1:1).

Supplementary Fig. 6 related to Fig. 6. GD2 expression following CASD1 silencing in MES cells.

Supplementary Fig. 7 related to Fig. 7. A. Long-term cytotoxicity (5 days) of UT (B) and GD2-ζCAR (C) Vδ2T cells against mixed 3D-tumoroids of RFP-GBM1:GFP-GBM10 (MES:PN).B-D. B-D. Short-term cytotoxicity (24 hours) of Vδ2T cell subsets against 2D-mixed cultures (B), RFP-MES cells (C) and GFP-PN cells (D).

## Material and methods

### Primary culture of human glioblastoma

All procedures involving human participants were in accordance with the ethical standards of the ethic National Research Committee and with the 1964 Helsinki Declaration and its later amendments or comparable ethical standards (Ref. MESR DC-2014–2206, institutional approval from committee patient protection board, “CPP Ouest IV, approval number “Dossier 06/15”). The patients or their guardians/legally authorized representatives/next of kin provided written informed consent for participation in the use of samples. Primary cultures of human GBM were obtained after the mechanical dissociation of surgical tumor samples from patients undergoing surgical intervention at the Department of Neurosurgery at “Centre Hospitalier Universitaire de Nantes” and the “Tumorothèque IRCNA” ^60^. Primary cultures were stored and frozen in liquid nitrogen in 50% medium/40% Fetal Calf Serum (FCS) (Biosera S00IK10004)/10% DMSO (Sigma-Aldrich D2660). Cells were kept in a wet atmosphere at 37°C and 5% CO_2_ for at most 2 months following defrosting. The neurobasal (NBL) medium was used for the culture of primary GBM, which is composed of DMEM/F12 (Gibco 21331-020) supplemented with 2 mM L-glutamine (Gibco 25030081), antibiotics (Streptomycin 100 µg/mL and penicillin 100 IU/mL, Gibco 15140-122), growth factors B27 (Gibco 17502-048), N2 (Gibco 17504-044), 25 ng/mL b-FGF (Miltenyi Biotec 130-093-838), 20 ng/mL EGF (Miltenyi Biotec 130-093-825) and 2 µg/mL heparin (Sigma-Aldrich H3149). Two third of the culture medium was renewed 3 times a week after cell centrifugation. As these cultures were proliferating as neurospheres, they were incubated with 400-600 IU/mL accutase (Sigma-Aldrich A6964) for at least 5 mins before any numeration to dissociate these spheres.

### Primary culture of human Vδ2T cells

Peripheral blood mononuclear cells (PBMC) were isolated from blood bags of consenting healthy donors from the Etablissement Français du Sang (Blood product transfer agreement relating to biomedical research protocol 97/5-B—DAF 03/4868). PBMCs from donors were isolated using density gradient centrifugation with lymphocyte separation medium (Eurobio Scientific, CMSMSL01-01), then frozen in FCS/10% DMSO and stored in liquid nitrogen.

Vδ2T cells were amplified from these PBMC (4.10^6^ in 1 well of a 24M G-Rex® (*Wilson Wolf #80192M*) in the presence of Zoledronate 5 µM (Zometa, AltanPharma) X-vivo15 medium (Lonza 02-060Q) supplemented with 4% pooled allogeneic human serum (Team 12 CRCI2NA), or RPMI 1640 medium (Gibco 31870) supplemented with 8% HS, 2 mM glutamax (Gibco 35050-038) and 100 μg/mL streptomycin and 100 IU/mL penicillin (Gibco 15140-122). At day 3 of the culture, 100 IU/mL of recombinant IL-2 (Proleukin Clinigen) was added. From day 6, Vδ2T cells were cultured with 100 IU/mL IL-2 and 100 U/ml IL-15 (Miltenyi Biotec 130-095-760). Vδ2T cells were used once returned to a quiescent state, between 18 and 35 days after the stimulation. For further amplification, Vδ2T cells were stimulated with irradiated (35 Gy), pooled allogeneic feeder cells (PBMC from 2 donors and three EBV-LCL), 5 µM zoledronate and 300 IU/ml recombinant IL-2, and amplified for 18 more days.

### Retroviral vector production and T-cell transduction

The 2^nd^ generation CAR-OGD2 and the CAR-GD2 scFv were derived from the variable part of the 8B6 or the 14G2a monoclonal antibodies respectively. These scFv were linked to a human mutated IgG1 as the spacer, CD28 as the costimulatory domain, and CD3ζ as the transducing domain. As a control, ΔCAR without CD3ζ as the transducing domain was used. The CAR constructs were synthesized by PCR (GeneCust Europe, Dudelange, Luxembourg) and cloned into a retroviral pMX vector. Stable Phoenix-Ampho packaging cells were selected and used to produce recombinant retroviral vectors. The phoenix-ampho-pMx-CAR were plated and after 48h the retroviral supernatant was collected, filtered through 0.45µm pore-size filters, and concentrated 30-fold using Retro-X Concentrator ® reagent (*TAKARA PT5063-2*) and stored at -80°C until use. Two days after the initial zoledronate stimulation, Vδ2T cells were transduced in the presence of 4 μg/mL polybrene (Millipore TR-1003-G). The culture medium was changed 24 hours after infection. Mock (non-transduced) controls were performed in parallel. The transduction efficiencies were assessed 9 days later by staining the CAR with an Alexa Fluor 647-conjugated F(ab’)2 fragment goat anti-mouse IgG F(ab’)2 fragment (Jackson Immunological Research 115-606-072) or Alexa Fluor 647-conjugated anti-G4S linker (Cell signaling 69782). Ten days after transduction, T cells were sorted by flow cytometry (BD AriaIII) on Vδ2 TCR and CAR expression.

### Inactivation of B4GALNT1 and CASD1 genes in GBM primary cultures

Inactivation of the genes coding for B4GALNT1 (B4GAL) or CASD1 was realized using CRISPR/Cas9 technology, after the transduction of lentiviral particles coding for the endonuclease Cas9 and a specific guide sequence. These particular sequences were 5’-CACCGCGTCCCGGGTGCTCGCGTAC-3’ for B4GALNT1, and 5′-ATGCAATATGTTTATCTACA-3′ for CASD1, based on the sequence published in Baumann et al 2015^61^. The plasmid was produced after the transformation and selection of Stbl3 competent bacteria (Invitrogen C737303) in Lysogeny Broth (LB) medium containing 100 µg/mL ampicillin (Sigma-Aldrich A-9518).

HEK293 cells were cultivated in DMEM medium (Gibco 21969035) supplemented with 5 % FCS, 2 mM L-glutamine (Gibco 25030081), and antibiotics. To produce viral particles, cells were transfected with the plasmids coding for the different constructions (CRISPR/Cas9 LCV2, KO B4GAL, or KO CASD1). Two other plasmids, MD2 (Addgene, plasmid #20864) and psPAX2 (Addgene, plasmid #12260), were also used to allow for the production of viral particles and packaging of the plasmid, respectively. These plasmids were incubated with HEK293T cells and lipofectamine 2000 (Invitrogen 11668019) for 6 hours at 37°C in Opti-MEM medium (Gibco 31985070), then the medium was changed to fresh DMEM/5% FCS, and cells were incubated for 48 hours at 37°C. The supernatant was collected, filtered, and added to GBM cells overnight. Cells were then washed and put back in the NBL medium. A selection of transduced cells was achieved by adding 2 µg/mL of puromycin over one week.

### GD2 and OGD2 cell surface staining

For GD2 and OGD2 staining, cells were first washed once with PBS and once with PBS/BSA 1% (Sigma-Aldrich A2153-100G). The cells were then incubated with 10 µg/mL of isotype (DOTA IgG2a or IgG3, OGD2 pharma), 14G2a (Bioxcell BE0318), or 8B6 (IgG3, OGD2 pharma) in PBS/BSA 1% for 30 mins at 4°C. After two additional washing steps in PBS/BSA 1%, cells were incubated with 3 µg/mL FITC-conjugated goat anti-mouse (Jackson Immunology AB_2338589) for 30-45 mins at 4°C. Cells were washed two additional times and kept in cold PBS before the acquisition with an Accuri C6 plus flow cytometer (BD Biosciences). Data were analyzed using Flowjo 10 software. The Ratio of Fluorescence Intensity (RFI) was determined using the geometric mean of fluorescence intensity measured in the marked population, reported to the intensity measured with the control isotype.

### Gangliosides extraction and purification

GBM1 and GBM15 cells were lyophilized and extracted twice with CHCl_3_/CH_3_OH (2:1, v/v) and once with CHCl_3_/CH_3_OH (1:2, v/v) using intermediary centrifugations at 2,500g for 20 min. Combined supernatants were dried under a nitrogen stream, subjected to mild saponification in 0.1 M NaOH in CHCl_3_/CH_3_OH (1:1, v/v) at 37 °C for 2 h, and evaporated to dryness. Samples were reconstituted in CH_3_OH/0.1% TFA in water (1:1, v/v) and applied to a reverse-phase C18 cartridge (Supelco). After washing with CH_3_OH/0.1% TFA in water (1:1, v/v), glycosphingolipds (GSLs) were eluted by CH_3_OH, CHCl_3_/CH_3_OH (1:1, v/v) and CHCl_3_/CH_3_OH (2:1, v/v). The elution fraction was dried under a nitrogen stream before structural analysis.

### Mass spectrometry analysis of gangliosides

Gangliosides were permethylated using the sodium hydroxide/DMSO slurry method^62^. Briefly, gangliosides were incubated with the permethylation reagent at room temperature for 2 hours under constant agitation. The reaction was quenched by the addition of water, and the permethylated glycans were extracted with chloroform (CHCl₃) and washed at least seven times with water to remove residual reagents. The permethylated gangliosides were analyzed using a MALDI-QIT-TOF mass spectrometer (Shimadzu AXIMA Resonance, Shimadzu Europe, Manchester, UK) in positive ion mode. For sample preparation, 1 μL of the permethylated GSL solution in CHCl₃ was mixed directly on the MALDI target with 1 μL of superDHB matrix (10 mg/mL in CHCl₃/CH₃OH, 1:1, v/v). Spectra were acquired using the “mid mode” over a mass range of m/z 1000–3000. The laser was operated at a power setting of 100, with two laser shots taken at each of the 200 distinct positions within the sample spot.

### Vδ2T cells degranulation assay

10^5^ GBM cells were first seeded in wells of a 96-well round-bottom plate in NBL medium in the presence or not of 20 µM zoledronate overnight at 37°C. The supernatant was then discarded and the T cells were added at a 1:1 effector:target (E:T) ratio, in RPMI medium containing 5 µM of monensin (Merck 5273-1G) and anti-CD107a-AF647 antibody (Biolegend 328612) and supplemented with 10 % FCS, 2 mM L-glutamine, and antibiotics. When indicated, GBM cultures were incubated with 10 µg/mL of a BTN3A blocking antibody (103.2, Innate Pharma) for 15 mins before adding Vδ2T cells. In some experiments, Vδ2 T cells were incubated alone with an anti-Vδ2 TCR (Biolegend 331406) antibody or the 1A7 antibody (Absolute antibody, Ab0227-10.0), at 10 μg/mL or 2.5 μg/mL, respectively. After 4h of coculture, the cells were washed and stained with the same anti-Vδ2 TCR antibody for 20 mins RT. After an additional wash step, the cells were analyzed by flow cytometry using an Accuri C6 plus flow cytometer (BD Biosciences). Data were then analyzed using Flowjo 10 software.

### Cytokines release assay

30,000 GBM cells were seeded per well of a 96-well round-bottom plate in NBL medium in the presence or not of 20 µM zoledronate overnight at 37°C. The next morning, the supernatant was removed, T cells were added at a 5:1 E:T ratio in RPMI/10% FCS medium, and the plate was left to incubate for 4 hours at 37°C. The cells were then centrifuged and the supernatant was recovered and frozen at -80°C. Cytokines were dosed using Legendplex CD8/NK panel (Biolegend 741187) following the manufacturer’s procedure. Briefly, samples and standards were diluted in the assay buffer and then incubated with a mix of thirteen fluorescent beads (APC), each specific for a different cytokine, for 2 hours at room temperature and under stirring. After two wash steps, biotinylated detection antibodies were added for 1 hour at room temperature and under stirring. Streptavidin-PE was then added for 30 minutes at room temperature and under stirring. After two wash steps, samples were read using an Attune Nxt flow cytometer (Thermofisher Scientific). Data obtained from these experiments were analyzed using the manufacturer’s software.

### Flow-based cytotoxicity assays

30,000 GBM cells were seeded per well of a 96-well round-bottom plate in NBL medium in the presence or not of 20 µM zoledronate overnight at 37°C. The next morning, the supernatant was removed, T cells were added at different E:T ratios (1:1, 5:1, or 10:1) in RPMI/10% FCS medium, and the plate was left to incubate for 24 hours at 37°C. The cells were transferred in a conical bottom plate, washed once in PBS, and incubated with the Zombie Viability dye (Biolegend 423113) for 20 mins at room temperature. The cells were washed once in PBS/BSA 1% and then stained with CD3-APC antibody (Biolegend 344812) for 20 mins at room temperature. After an additional wash in PBS/BSA 1%, samples were acquired using an Attune Nxt flow cytometer. During the analysis, after the elimination of the doublets and the debris, CD3-positive events were excluded to assess only the percentage of dead tumor cells. Flow cytometry data were analyzed using Flowjo 10 software. The percentage of cytotoxicity was calculated based on the following formula: % cytotoxicity = 100*(% dead cells in sample - % dead cells alone) / (100 - % dead cells alone).

### Microscopy-based cytotoxicity assays

For 3D killing assays, a total of 6000 fluorescent GBM cells were seeded per well of a 96-well round-bottom ultra-low attachment plate (Corning 7007) in NBL medium, using only one subtype or mixing both subtypes with a 1:1 ratio for mixed tumoroids. They were then centrifuged at 270G for 3 mins and kept at 37°C for 3 days to let the cells form spheres. Three-quarters of the medium was removed and Vδ2T cells were added in a 1:1, 5:1, or 10:1 E:T ratio in NBL medium supplemented with 100 IU/mL recombinant IL-2. In some wells, 20 µM of zoledronate was added to assess the maximum effect of the Vδ2 T cells. Fluorescent cells were then followed for 5 days after adding T cells using the Incucyte (Sartorius) or Cytation (Agilent) instrument. Mean fluorescence intensity and area of fluorescence were both quantified for the GFP and the RFP channels, using the Incucyte software or ImageJ. The integrated intensity of fluorescence was calculated as the product of those two parameters, and an estimation of the lysis caused by Vδ2T cells was calculated as follows: % lysis = 100*(1 - (Fluorescence_Sample_ – Fluorescence_Zoledronate_) / (Fluorescence_Ctrl_ – Fluorescence_Zoledronate_).

### Synapse visualization by confocal microscopy

The coverslips were added in plate 24 wells with poly L-Lysine (Sigma, P4707) for 2 hours at 37°C. After, they were washed 3 times with PBS. The primary GBM cells expressing RFP-like protein were added at 10^5^ cells per well and incubated overnight at 37°C. 10^5^ Vδ2T cells were added per well for 15 mins at room temperature and 15 mins at 37°C. The coverslips were washed with PBS and then incubated with an anti-TCR Vδ2-FITC mAb (Biolegend, 331406) on ice at 4°C for 1 hour. After one PBS/BSA 1% wash, coverslips were incubated with antibodies anti-G4S linker AlexaFluor 647 (CellSignaling, 69782S) and antibody donkey anti-mouse IgG AlexaFluor 488 (ThermoFischer Scientific, A21202) on ice at 4°C for 1 hour. Once washed with PBS/BSA 1%, then PBS, cells were fixed with 4% paraformaldehyde (Sigma-Aldrich 441244) at room temperature for 15 mins. Nuclei were counterstained with Hoechst (Interchim, FP-BB1340) at room temperature for 15 mins. The coverslips were mounted using Fluoromount-G™ Mounting Medium (Interchim, FP-483331).

### Immunohistochemistry and immunofluorescence analyses on primary GBM samples

Primary GBM samples came from patients with IDH-wildtype GBM included in the French GB biobank (FGB)^37^. FGB was declared to the French Ministry of Health and Research (declaration number: DC-2011-1467, cession authorization number: AC-2023-5473, BRIF (bioresource research impact factor) number: BB-0033-00093). The protocols and regulations of the FGB were approved by the CPP OUEST II ethics committee (CB 2012/02, date of approval: 20 December 2011) and the CNIL (“Commission Nationale de l’Informatique et des Libertés”, the French national data protection authority, no. 1476342, date of approval: 10 October 2011). All adult patients included in this study signed an informed consent form for the inclusion of their clinical data and samples in the FGB. Immunohistochemistry was performed for NKG2D (ULBP2/5/6; ULBP3; MICA/B) and TCR Vδ2 (BTN2A1) ligands on 4 μm sections of FFPE tissue blocks from 15 primary GBM samples with an automated Leica BOND III (Leica Biosystems, Nanterre, France) following the manufacturer’s instructions. Antibodies were diluted in BOND Primary Antibody Diluent (Leica) as follows: anti-ULBP2/5/6 antibody (Bio-Techne, Rennes, France; 1.5 µg/mL), anti-ULBP3 antibody (CliniSciences, Nanterre, France; 3 µg/mL), anti-MICA/B (Bio-Techne; 25 µg/mL), and BTN2A1 (Life Technologies, Courtaboeuf, France; 1.5 µg/mL). EDTA-based pH 9 was used as an antigen retrieval solution. Primary antibody binding to GBM tissue sections was visualized with BOND Polymer Refine Detection (Leica Biosystems). A colon brain metastasis tissue was used as a positive control for immunostaining. Digital images were captured with an Aperio CS2 scanner (Leica Biosystems) fitted with a 20x objective and analyzed with Aperio ImageScope software v12.3.2.8009 (Leica Biosystems). A semi-quantitative analysis of IHC images was performed with Fiji software as described by Crowe and Yue (2019). The percent positivity was calculated as the number of positive pixels divided by the total number of pixels in the area analyzed, multiplied by 100. Six tumor areas of identical size were analyzed in each section and the mean percent positivity was calculated. Immunofluorescence (IF) was performed for GD2 and OGD2 on 8 μm frozen tissue sections from 12 primary GBM samples. Sections were treated with ice-cold acetone, rehydrated with PBS, and blocked in 10% human serum in BOND Primary Antibody Diluent (Leica Biosystems). Anti-GD2 mAb 14G2a (5 µg/mL, Becton Dickinson, Pont de Claix, France) and anti-OGD2 mAb 8B6 (10 µg/mL) were added onto the sections overnight, at 4°C. Antibodies were detected with secondary antibodies conjugated to Alexa Fluor 488 (Fisher Scientific, Illkirch, France). Nuclei were counterstained with DAPI (Sigma-Aldrich Chimie, Saint Quentin Fallavier, France). Mouse IgG2a and mouse IgG3 were used as negative controls. Sections were observed under a fluorescence Axioscope 2 optical microscope (Zeiss, Le Pecq, Germany) and analyzed with Zen 3.2 (blue edition) software (Zeiss). The mean percent positivity of tumor areas was calculated as described above.

### Transcriptomic analysis

Primary GBM cells were washed twice in PBS, and total RNA was extracted using the RNeasy Mini Kit (Qiagen) with on-column DNase I treatment, following the manufacturer’s instructions. RNA quantity and quality were assessed using a NanoDrop ND-1000 spectrophotometer (Thermo Fisher Scientific) and an Agilent 2100 Bioanalyzer (Agilent Technologies), respectively. For RNA sequencing, samples were processed using 3′ SRP RNA-seq. Gene expression normalization and differential gene expression analysis were performed using the DESeq2 package (Bioconductor) in R. The dataset was normalized against a virtual median sample, and low-count genes (<10 reads across all samples) were excluded ^63^. Data were then subjected to variance-stabilizing transformation prior to downstream analyses. Gene Set Enrichment Analysis (GSEA) was carried out using the Broad Institute’s GSEA software ^64^ and RStudio version 4.1.1 ^65,66^. Annotated gene sets from the Molecular Signatures Database (MSigDB) were used for ganglioside and ceramide metabolism pathways. Gene sets defined by Neftel et al. were employed to assess glioblastoma cell states ^5^. Samples were grouped according to molecular subtype as previously described, and permutations were conducted based on gene sets. RNA-seq data will be deposited in the Gene Expression Omnibus (GEO) (submission in progress).

### Statistical analysis

Unless otherwise specified, data are presented as mean ± standard deviation (s.d.) and were analyzed using GraphPad Prism version 10 (GraphPad Software, Inc.). Two-way ANOVA followed by appropriate multiple comparisons tests was used depending on the assay: Tukey’s multiple comparisons test for CD107a degranulation and Zombie-based lysis assays; Sidak’s multiple comparisons test for ganglioside (GD2, OAcGD2) expression and cytokine production; and Dunnett’s multiple comparisons test for tumoroid lysis assays. For in vivo experiments, survival curves were analyzed using the log-rank (Mantel-Cox) test with p-values adjusted with the Bonferroni method for multiple group comparison. Statistical significance was defined as follows: P < 0.05 (*), P < 0.01 (**), P < 0.001 (***), and P < 0.0001 (****). Symbols indicating group comparisons are as follows: * indicates significance compared to untransduced (UT) cells; $ indicates significance between paired ΔCAR and CAR T cells; and # indicates significance between GD2-CAR and OGD2-CAR T cells.

Each dot in the graphs represents an independent biological replicate. No statistical methods were used to predetermine sample sizes but we used adequate numbers of biological replicates that would provide statistically significant results on the basis of our previous experience. In vivo experiments were randomized across treatment groups. The investigators were not blinded to group allocation during experimental procedures but were blinded to outcome assessments.

## REFERENCES

1. Stupp, R. et al. Radiotherapy plus concomitant and adjuvant temozolomide for glioblastoma. N. Engl. J. Med. 352, 987–996 (2005).

2. Stupp, R. et al. Effects of radiotherapy with concomitant and adjuvant temozolomide versus radiotherapy alone on survival in glioblastoma in a randomised phase III study: 5-year analysis of the EORTC-NCIC trial. Lancet Oncol. 10, 459–466 (2009).

3. Louis, D. N. et al. The 2021 WHO Classification of Tumors of the Central Nervous System: a summary. Neuro Oncol 23, 1231–1251 (2021).

4. Verhaak, R. G. W. Moving the needle: Optimizing classification for glioma. Science Translational Medicine 8, 350fs14–350fs14 (2016).

5. Neftel, C. et al. An Integrative Model of Cellular States, Plasticity, and Genetics for Glioblastoma. Cell 178, 835–849.e21 (2019).

6. Wang, L. et al. A single-cell atlas of glioblastoma evolution under therapy reveals cell-intrinsic and cell-extrinsic therapeutic targets. Nat Cancer 3, 1534–1552 (2022).

7. Oizel, K. et al. Efficient Mitochondrial Glutamine Targeting Prevails Over Glioblastoma Metabolic Plasticity. Clin. Cancer Res. 23, 6292–6304 (2017).

8. Mao, P. et al. Mesenchymal glioma stem cells are maintained by activated glycolytic metabolism involving aldehyde dehydrogenase 1A3. Proc. Natl. Acad. Sci. U.S.A. 110, 8644–8649 (2013).

9. Halliday, J. et al. In vivo radiation response of proneural glioma characterized by protective p53 transcriptional program and proneural-mesenchymal shift. Proc. Natl. Acad. Sci. U.S.A. 111, 5248–5253 (2014).

10. Grossman, S. A. et al. Immunosuppression in patients with high-grade gliomas treated with radiation and temozolomide. Clin Cancer Res 17, 5473–5480 (2011).

11. Akhavan, D. et al. CAR T cells for brain tumors: Lessons learned and road ahead. Immunol Rev 290, 60–84 (2019).

12. Meyer, M. et al. Single cell-derived clonal analysis of human glioblastoma links functional and genomic heterogeneity. Proc Natl Acad Sci U S A 112, 851–856 (2015).

13. Renoult, O. et al. Metabolic profiling of glioblastoma stem cells reveals pyruvate carboxylase as a critical survival factor and potential therapeutic target. Neuro Oncol 26, 1572–1586 (2024).

14. Segerman, A. et al. Clonal Variation in Drug and Radiation Response among Glioma-Initiating Cells Is Linked to Proneural-Mesenchymal Transition. Cell Rep 17, 2994–3009 (2016).

15. Schmitt, M. J. et al. Phenotypic mapping of pathological crosstalk between glioblastoma and innate immune cells by synthetic genetic tracing. Cancer Discov 11, 754–777 (2021).

16. Patel, A. P. et al. Single-cell RNA-seq highlights intratumoral heterogeneity in primary glioblastoma. Science 344, 1396–1401 (2014).

17. Lin, H. et al. Understanding the immunosuppressive microenvironment of glioma: mechanistic insights and clinical perspectives. Journal of Hematology & Oncology 17, 31 (2024).

18. Brown, C. E. et al. Bioactivity and Safety of IL13Rα2-Redirected Chimeric Antigen Receptor CD8+ T Cells in Patients with Recurrent Glioblastoma. Clin Cancer Res 21, 4062–4072 (2015).

19. Krenciute, G. et al. Characterization and Functional Analysis of scFv-based Chimeric Antigen Receptors to Redirect T Cells to IL13Rα2-positive Glioma. Mol Ther 24, 354–363 (2016).

20. O’Rourke, D. M. et al. A single dose of peripherally infused EGFRvIII-directed CAR T cells mediates antigen loss and induces adaptive resistance in patients with recurrent glioblastoma. Sci Transl Med 9, eaaa0984 (2017).

21. Schmidts, A. et al. Tandem chimeric antigen receptor (CAR) T cells targeting EGFRvIII and IL-13Rα2 are effective against heterogeneous glioblastoma. Neurooncol Adv 5, vdac185 (2022).

22. Ahmed, N. et al. HER2-specific T cells target primary glioblastoma stem cells and induce regression of autologous experimental tumors. Clin Cancer Res 16, 474–485 (2010).

23. Thomas, P., Paris, P. & Pecqueur, C. Arming Vδ2 T cells with chimeric antigen receptors to combat cancer. Clin Cancer Res https://doi.org/10.1158/1078-0432.CCR-23-3495 (2024) doi:10.1158/1078-0432.CCR-23-3495.

24. Chauvin, C. et al. NKG2D Controls Natural Reactivity of Vγ9Vδ2 T Lymphocytes against Mesenchymal Glioblastoma Cells. Clin Cancer Res 25, 7218–7228 (2019).

25. Wilhelm, M. et al. Gammadelta T cells for immune therapy of patients with lymphoid malignancies. Blood 102, 200–206 (2003).

26. Sebestyen, Z., Prinz, I., Déchanet-Merville, J., Silva-Santos, B. & Kuball, J. Translating gammadelta (γδ) T cells and their receptors into cancer cell therapies. Nat Rev Drug Discov 19, 169–184 (2020).

27. Alvarez-Rueda, N. et al. A Monoclonal Antibody to O-Acetyl-GD2 Ganglioside and Not to GD2 Shows Potent Anti-Tumor Activity without Peripheral Nervous System Cross-Reactivity. PLoS One 6, (2011).

28. Mount, C. W. et al. Potent antitumor efficacy of anti-GD2 CAR T-cells in H3K27M+ diffuse midline gliomas. Nat Med 24, 572–579 (2018).

29. Majzner, R. G. et al. GD2-CAR T cell therapy for H3K27M-mutated diffuse midline gliomas. Nature 603, 934–941 (2022).

30. Liu, Z. et al. Safety and antitumor activity of GD2-Specific 4SCAR-T cells in patients with glioblastoma. Mol Cancer 22, 3 (2023).

31. Yu, L. et al. GD2-specific chimeric antigen receptor-modified T cells for the treatment of refractory and/or recurrent neuroblastoma in pediatric patients. J Cancer Res Clin Oncol 148, 2643–2652 (2022).

32. Straathof, K. et al. Antitumor activity without on-target off-tumor toxicity of GD2–chimeric antigen receptor T cells in patients with neuroblastoma. Sci. Transl. Med. 12, eabd6169 (2020).

33. Heczey, A. et al. CAR T Cells Administered in Combination with Lymphodepletion and PD-1 Inhibition to Patients with Neuroblastoma. Mol Ther 25, 2214–2224 (2017).

34. Bufalo, F. D. et al. GD2-CART01 for Relapsed or Refractory High-Risk Neuroblastoma. New England Journal of Medicine 388, 1284–1295 (2023).

35. Ciccone, R. et al. GD2-Targeting CAR T-cell Therapy for Patients with GD2+ Medulloblastoma. Clinical Cancer Research 30, 2545–2557 (2024).

36. Locatelli, F. et al. GD2-targeting CAR T cells in high-risk neuroblastoma: a phase 1/2 trial. Nat Med 31, 3689–3699 (2025).

37. Clavreul, A. et al. The French glioblastoma biobank (FGB): a national clinicobiological database. J Transl Med 17, 133 (2019).

38. Faubert, B., Solmonson, A. & DeBerardinis, R. J. Metabolic reprogramming and cancer progression. Science 368, (2020).

39. Mabe, N. W. et al. Transition to a mesenchymal state in neuroblastoma confers resistance to anti-GD2 antibody via reduced expression of ST8SIA1. Nat Cancer 3, 976–993 (2022).

40. Prapa, M. et al. GD2 CAR T cells against human glioblastoma. *npj Precis*. Onc. 5, 1–14 (2021).

41. Cavdarli, S., Delannoy, P. & Groux-Degroote, S. O-acetylated Gangliosides as Targets for Cancer Immunotherapy. Cells 9, (2020).

42. Raghavan, S. et al. Microenvironment drives cell state, plasticity, and drug response in pancreatic cancer. Cell 184, 6119–6137.e26 (2021).

43. Gargett, T. et al. GD2-targeting CAR-T cells enhanced by transgenic IL-15 expression are an effective and clinically feasible therapy for glioblastoma. J Immunother Cancer 10, e005187 (2022).

44. de Billy, E. et al. Dual IGF1R/IR inhibitors in combination with GD2-CAR T-cells display a potent anti-tumor activity in diffuse midline glioma H3K27M-mutant. Neuro-Oncology 24, 1150–1163 (2022).

45. Tirosh, I. & Suvà, M. L. Tackling the Many Facets of Glioblastoma Heterogeneity. Cell Stem Cell 26, 303–304 (2020).

46. Majzner, R. G. et al. Tuning the Antigen Density Requirement for CAR T-cell Activity. Cancer Discov 10, 702–723 (2020).

47. Wang, Q. et al. Tumor evolution of glioma intrinsic gene expression subtype associates with immunological changes in the microenvironment. Cancer Cell 32, 42–56.e6 (2017).

48. Weiss, T. et al. NKG2D-Dependent Antitumor Effects of Chemotherapy and Radiotherapy against Glioblastoma. Clin Cancer Res 24, 882–895 (2018).

49. Weiss, T., Weller, M., Guckenberger, M., Sentman, C. L. & Roth, P. NKG2D-Based CAR T Cells and Radiotherapy Exert Synergistic Efficacy in Glioblastoma. Cancer Res 78, 1031–1043 (2018).

50. Baugh, R. et al. Targeting NKG2D ligands in glioblastoma with a bispecific T-cell engager is augmented with conventional therapy and enhances oncolytic virotherapy of glioma stem-like cells. J Immunother Cancer 12, e008460 (2024).

51. Baumeister, S. H. et al. Phase I Trial of Autologous CAR T Cells Targeting NKG2D Ligands in Patients with AML/MDS and Multiple Myeloma. Cancer Immunol Res 7, 100–112 (2019).

52. Jones, A. B., Rocco, A., Lamb, L. S., Friedman, G. K. & Hjelmeland, A. B. Regulation of NKG2D Stress Ligands and Its Relevance in Cancer Progression. Cancers (Basel) 14, 2339 (2022).

53. Ayala Ceja, M., Khericha, M., Harris, C. M., Puig-Saus, C. & Chen, Y. Y. CAR-T cell manufacturing: Major process parameters and next-generation strategies. J Exp Med 221, e20230903 (2024).

54. Straetemans, T. et al. GMP-Grade Manufacturing of T Cells Engineered to Express a Defined γδTCR. Front Immunol 9, 1062 (2018).

55. Zhang, T. et al. Clinical Safety and Efficacy of Locoregional Therapy Combined with Adoptive Transfer of Allogeneic γδ T Cells for Advanced Hepatocellular Carcinoma and Intrahepatic Cholangiocarcinoma. J Vasc Interv Radiol 33, 19–27.e3 (2022).

56. Vydra, J. et al. A Phase I Trial of Allogeneic γδ T Lymphocytes From Haploidentical Donors in Patients With Refractory or Relapsed Acute Myeloid Leukemia. Clinical Lymphoma Myeloma and Leukemia 23, e232–e239 (2023).

57. Neelapu, S. S. et al. A phase 1 study of ADI-001: Anti-CD20 CAR-engineered allogeneic gamma delta (γδ) T cells in adults with B-cell malignancies. JCO 40, 7509–7509 (2022).

58. Kobayashi, H. et al. Safety profile and anti-tumor effects of adoptive immunotherapy using gamma-delta T cells against advanced renal cell carcinoma: a pilot study. Cancer Immunol Immunother 56, 469–476 (2007).

59. Bennouna, J. et al. Phase I study of bromohydrin pyrophosphate (BrHPP, IPH 1101), a Vgamma9Vdelta2 T lymphocyte agonist in patients with solid tumors. Cancer Immunol Immunother 59, 1521–1530 (2010).

60. Salaud, C. et al. Mitochondria transfer from tumor-activated stromal cells (TASC) to primary Glioblastoma cells. Biochem Biophys Res Commun 533, 139–147 (2020).

61. Baumann, A.-M. T. et al. 9-O-Acetylation of sialic acids is catalysed by CASD1 via a covalent acetyl-enzyme intermediate. Nat Commun 6, 7673 (2015).

62. Groux-Degroote, S., Cavdarli, S., Yamakawa, N., Guérardel, Y. & Delannoy, P. Recent progress in O-acetylated gangliosides analysis and functions in cancer. https://doi.org/10.1039/9781839164538-00537 (2021) doi:10.1039/9781839164538-00537.

63. Love, M. I., Huber, W. & Anders, S. Moderated estimation of fold change and dispersion for RNA-seq data with DESeq2. Genome Biology 15, 550 (2014).

64. Downloads. https://www.gsea-msigdb.org/gsea/downloads.jsp.

65. Subramanian, A. et al. Gene set enrichment analysis: A knowledge-based approach for interpreting genome-wide expression profiles. Proceedings of the National Academy of Sciences 102, 15545–15550 (2005).

66. Mootha, V. K. et al. PGC-1α-responsive genes involved in oxidative phosphorylation are coordinately downregulated in human diabetes. Nat Genet 34, 267–273 (2003).

