## Supplemental Figures for "Functional diversity of Vγ9Vδ2 T cells overcomes glioblastoma state plasticity and antigen heterogeneity"

Supp data related to Figure 1

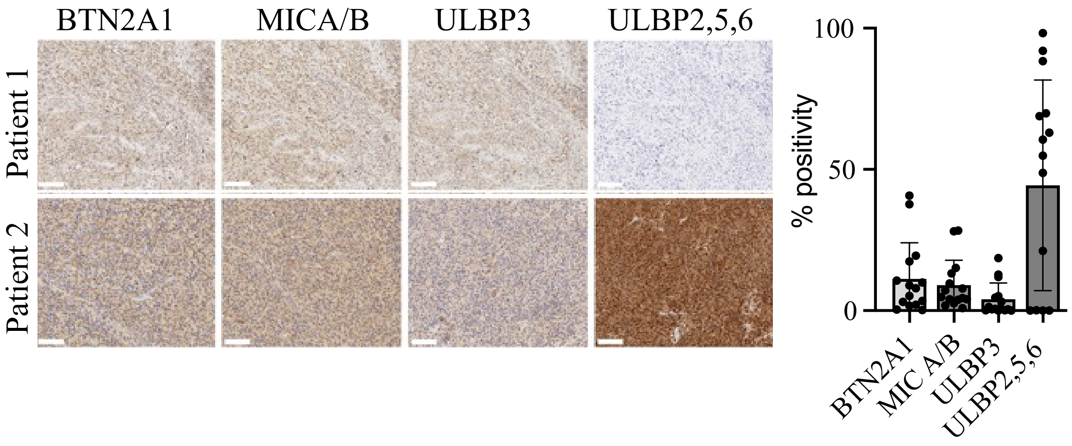

Supp data related to Figure 2

A

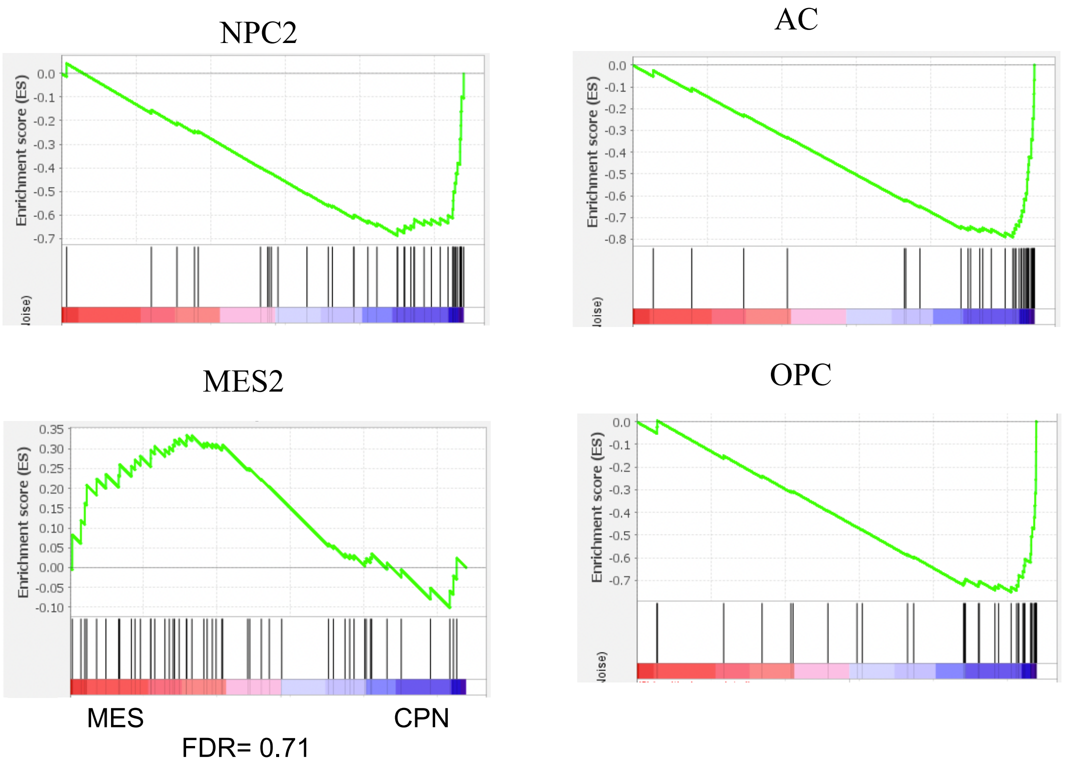


B

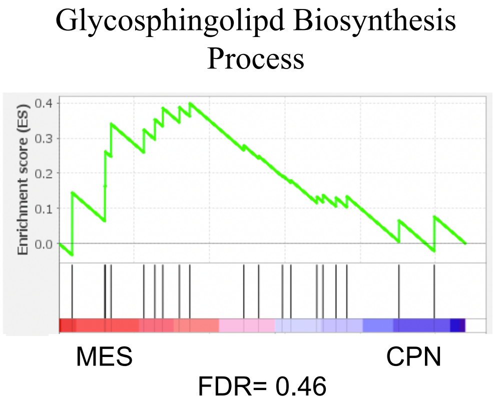


C

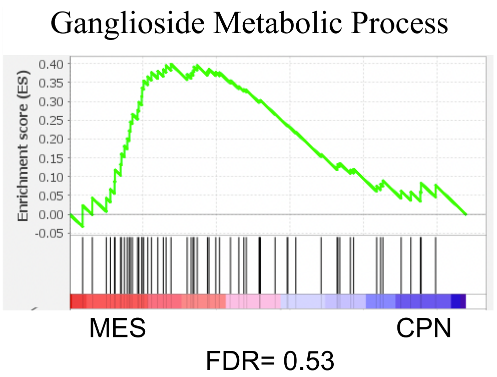


D

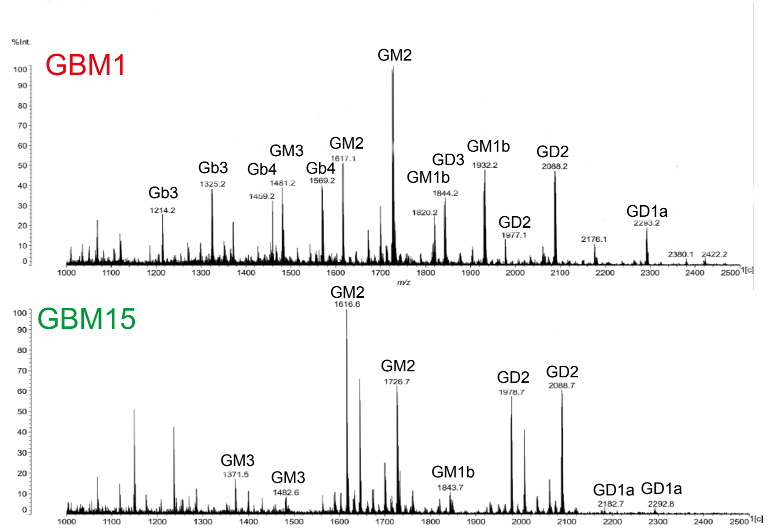

Supp data related to Figure 3

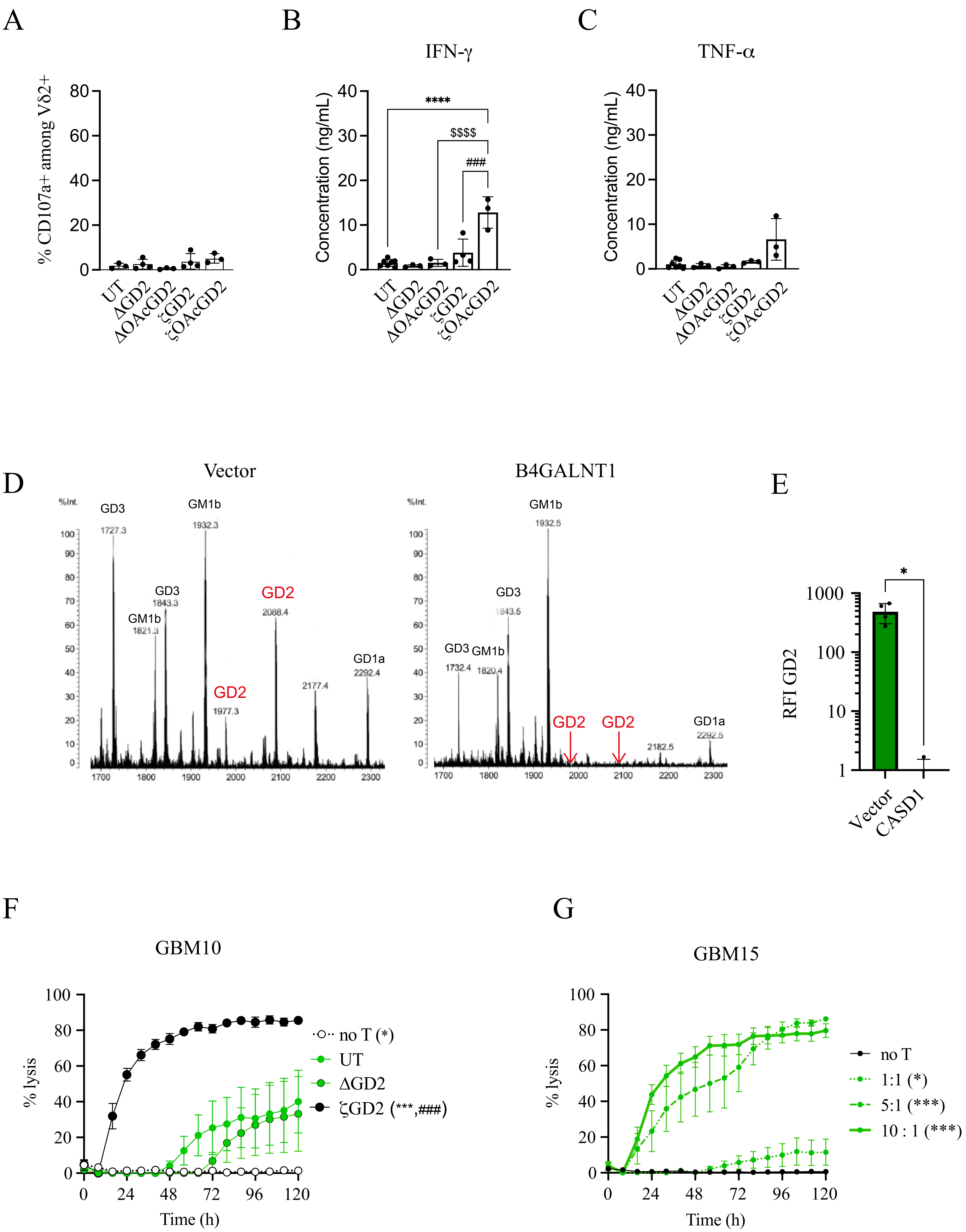

Supp data related to Figure 4

A

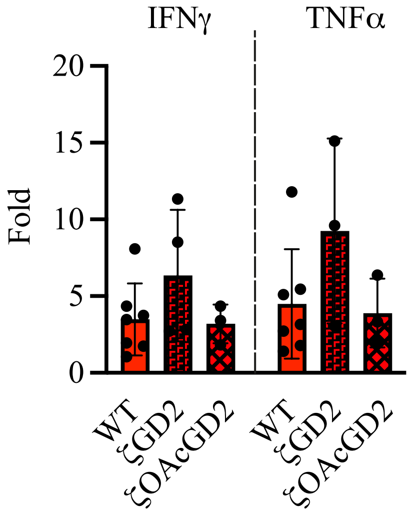


B

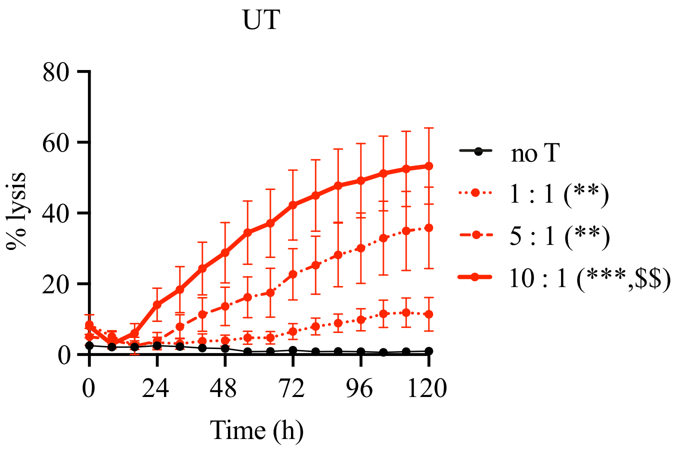


C

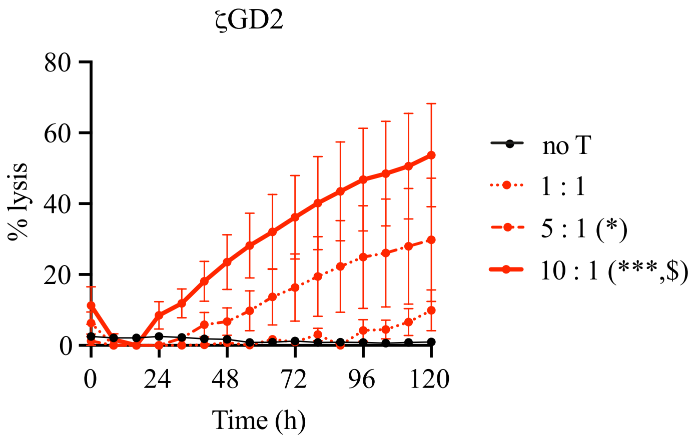

Supp data related to Figure 5

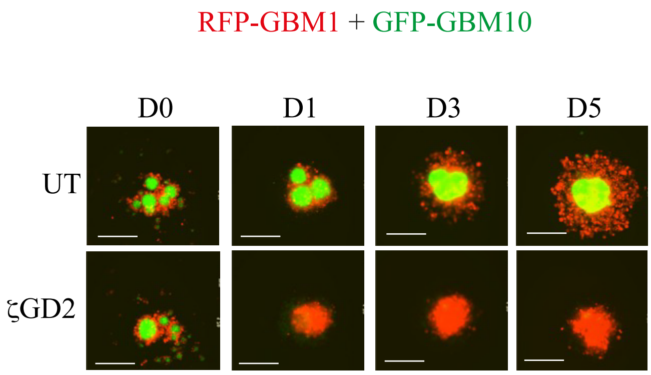

Supp data related to Figure 6

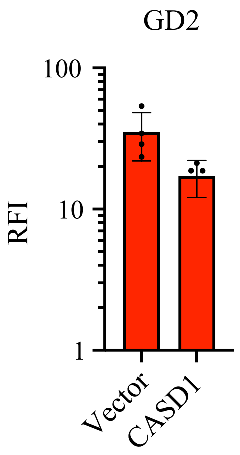

Supp data related to Figure 7

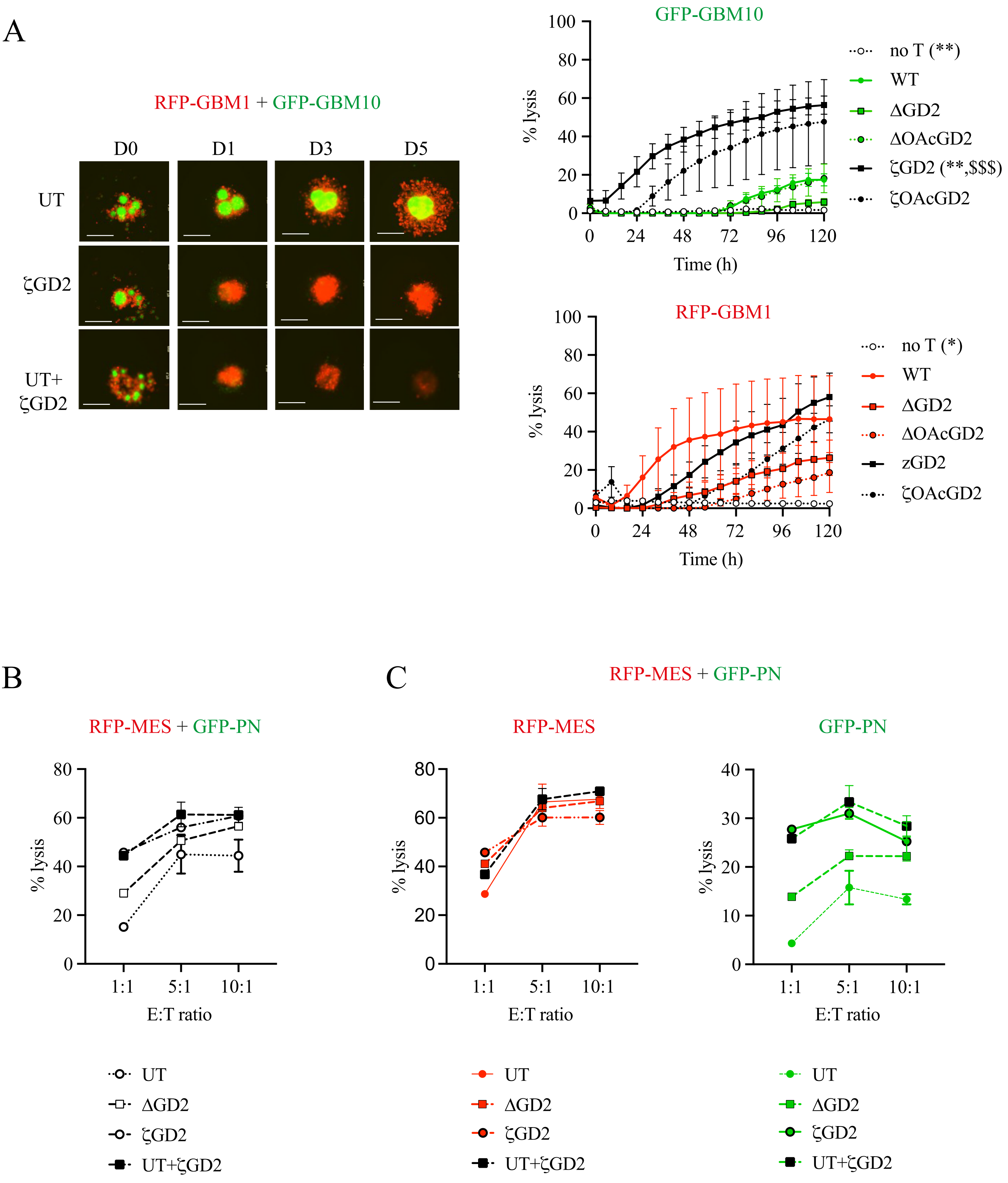
